# Proteomic Characterization of Intrahepatic Cholangiocarcinoma Identifies Distinct Subgroups and Proteins Associated with Time-To-Recurrence

**DOI:** 10.1101/2024.02.28.582093

**Authors:** Tilman Werner, Klara-Luisa Budau, Miguel Cosenza-Contreras, Frank Hause, Konrad Kurowski, Niko Pinter, Julia Schüler, Martin Werner, Carlie Sigel, Laura H. Tang, Peter Bronsert, Oliver Schilling

## Abstract

**Background & Aims:** Intrahepatic cholangiocarcinoma (ICC) is a poorly understood cancer with dismal survival and high recurrence rates. ICCs are often detected in advanced stages. Surgical resection is the most important first-line treatment but limited to non-advanced cases, whereas chemotherapy provides only a moderate benefit. The proteome biology of ICC has only been scarcely studied and the prognostic value of initial ICC’s proteomic features for the time-to-recurrence (TTR) remains unclear.

**Methods:** We dissected formalin-fixed, paraffin-embedded samples from 80 tumor– and 77 matching adjacent non-malignant (TANM) tissues. All samples were measured via liquid-chromatography mass-spectrometry (LC-MS/MS) in data independent acquisition mode (DIA).

**Results:** Tumor– and TANM tissue showed strongly different biologies and DNA-repair, translation, and matrisomal processes were upregulated in ICC. In a hierarchical clustering analysis, we determined two proteomic subgroups of ICC, which showed significantly diverging TTRs. Cluster 1, which is associated with a beneficial prognosis, was enriched for matrisomal processes and proteolytic processing, while cluster 2 showed increased RNA and protein turnover. In a second, independent Cox’ proportional hazards model analysis, we identified individual proteins whose expression correlates with TTR distribution. Proteins with a positive hazard ratio were mainly involved in carbon/glucose metabolism and protein turnover. Conversely, proteins associated with a low hazard ratio were mostly linked to the extracellular matrix. Additional proteome profiling of patient-derived xenograft tumor models of ICC successfully distinguished tumor and stromal proteins and provided insights into cell-matrix interactions.

**Conclusions:** We successfully determine the proteome biology of ICC and present two proteome clusters in ICC patients with significantly different TTR rates and distinct biological motifs. A xenograft model confirmed the importance of tumor-stroma interactions for this cancer.

## 1 Introduction

Cholangiocarcinomas (CCA) are the second most prevalent primary hepatic tumors and comprise 15% of all liver malignancies. According to their location in the liver bile ducts, CCAs are categorized as intrahepatic (ICC), perihilar or distal adenocarcinomas. CCAs possess different genetic, histologic, and clinical features but all are characterized by a late onset with diffuse clinical symptoms, frequent recurrences and dismal overall survival ^1–3^.

ICCs are rare cancers with rising incidence rates worldwide ^3^. Risk factors include stasis, parasitic or viral infections, alcohol and tobacco abuse, or obesity and diabetes ^4–6^. Higher case numbers in southeastern Asia can be explained by the prevalence of liver fluke infections^7^. Nevertheless, about 50% of cases occur without known risk ^8^.

ICCs grow in small bile ducts and ductules inside the liver, which makes them hard to access for histological probing. Diagnosis is further complicated by unspecific symptoms, imprecise imaging criteria and a lack of reliable non-invasive tumor markers ^1^. Therefore, most ICC patients present with large tumors, making extensive surgical resection the first-line treatment option ^9,10^. Although up to 70% of patients develop recurrences ^11^, no standardized treatment guidelines for adjuvant chemo-, targeted-, or immunologic therapies exist ^8,12–14^. In addition, an overall advantage of adjuvant chemo– or radiotherapy has not been confirmed ^10,15^. To enable successful tumor re-resections and personalized therapies, estimations of the risk for recurrence combined with cancer subtype classification are needed.

Pathological assessment of resected ICC largely stays confined to the criteria of the American Joint Committee on Cancer (AJCC) and the Union for International Cancer Control (UICC) ^1,10,15–18^. Frequently used classification into large– and small duct-type tumors was linked to different mutation profiles but is based on coarse definitions of subtypes and nomenclatures ^16^. In a previous study, we identified lymph node– and lymphangio-invasion, AJCC tumor staging, and tumor-budding as potential prognostic markers for the cohort presented here ^19^. More precise histopathological markers to assess tumor aggressiveness, enable tumor subtyping, and predict patients’ risk for early recurrence are lacking (Fig.1A).

**Figure 1.**
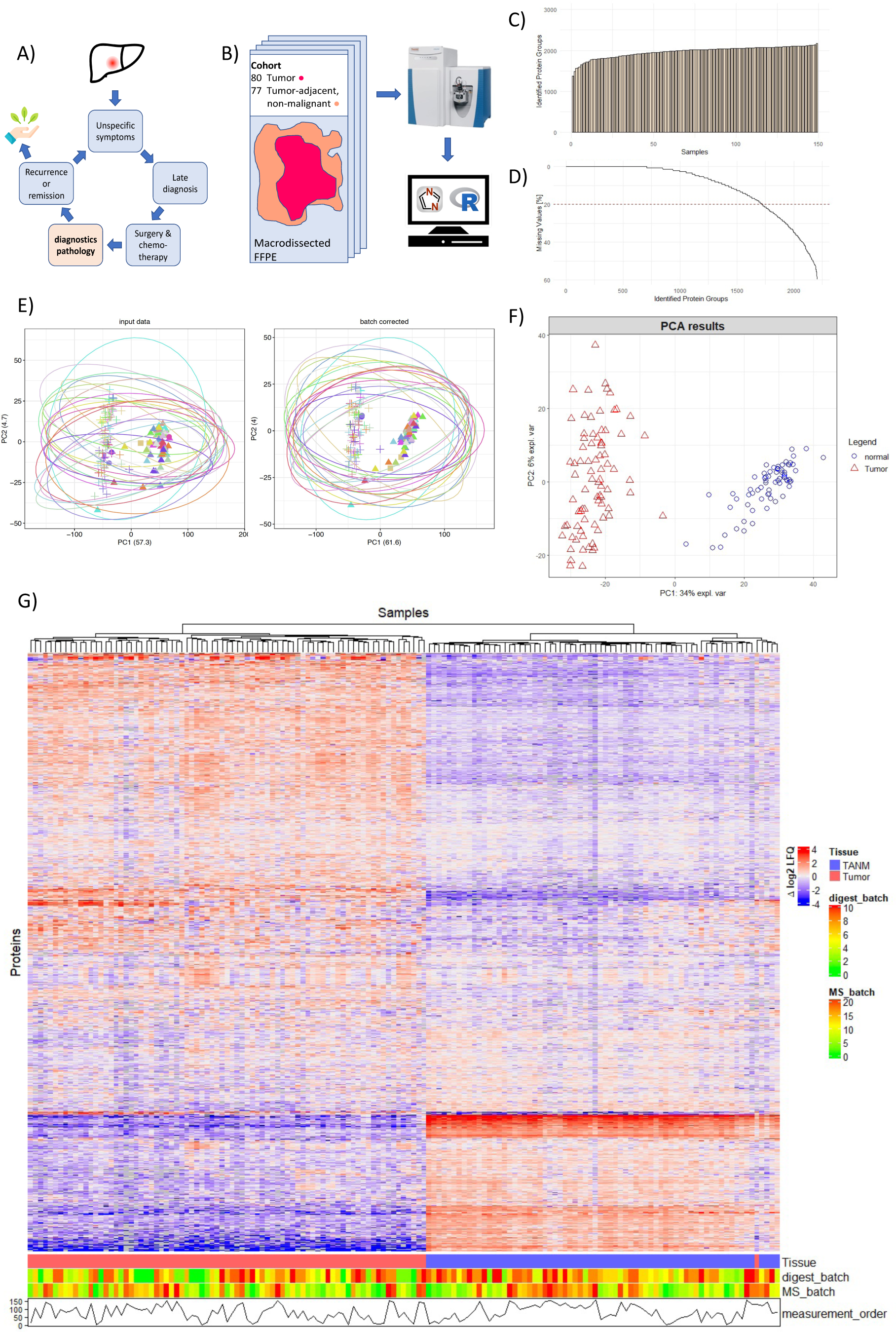
– cohort overview: A) Unspecific symptoms, late diagnoses and a lack of prognostic markers complicate treatment of ICC. B) Tumor and tumor-adjacent, non-malignant (TANM) tissue from 80 patients was macrodissected, measured, and analyzed. C) Protein-ID numbers across all samples of the cohort. D) Percentage of missing values across all detected proteins. E) PCAs comparing batch effects before and after ComBat batch correction. F) PCA of the entire cohort. G) Heatmap comparing the hierarchically clustered cohort to tissue type, digestion batch, measurement batch and measurement order.

Due to the rarity of ICC, this cancer remains insufficiently described on a molecular level and comprehensive omics-based studies are scarce ^1,3^. Genomic and transcriptomic studies have described the importance of *FGFR* fusions and *IDH1, ARID1A, BAP1, KRAS*, and *TP53* mutations for ICC genesis. Additionally, several publications proposed different molecular classifications into fibrotic, immune deserted, or innate– and acquired immunity-mediated inflammatory subtypes on genomic and transcriptomic level ^20–25^. A first integrated proteogenomic profile elaborating on ICC subtypes was presented by Dong et al ^26^ and followed by subsequent studies on the same cohort ^27^.

In recent years, there has been increasing interest in the characterization of tumor entities at the proteome level, which was further stimulated by groundbreaking studies from the Clinical Proteomic Tumor Analysis Consortium (CPTAC) ^28–30^. Since transcriptome alterations are only partially reflected on the protein level^31^ and cannot represent e.g. proteolytic events or post-translational modifications, proteomic profiling has become an integral component in understanding cancer biology. Advancing methods such as data-independent acquisition (DIA) have further facilitated cohort-wide studies with deep proteome coverages ^32^.

Here we present the proteomic characterization of 80 ICC tumors and 67 patient-matched, non-malignant adjacent tissues (TANM) highlighting tumor-specific proteome alterations. We identified two predominant proteomic subtypes of ICC which are strongly associated with time-to-recurrence. Further characterizations of proteolytic activity and a xenograft model emphasize the importance of tumor stroma interactions in ICC progression.

## 2 Materials and Methods

### 2.1 Ethics Statement

ICC tumors for this study were collected in the Memorial Sloan Kettering Cancer Center New York (MSKCC), USA, between 2009 and 2018. Written informed consent was obtained from all patients before inclusion. The study was conducted according to the Declaration of Helsinki. The MSKCC institutional review board approved the study (protocol number 16-1683A(3)).

### 2.2 ICC Cohort

All tissue specimens had been formalin-fixed, paraffin-embedded (FFPE), and were sliced to 10 µm cuts in the Department of Pathology at MSKCC. All histological slides were evaluated by two experienced pathologists at MSKCC according to the World Health Organization classification ^33^. TNM classifications were given based on the 8th edition of American Joint Committee on Cancer (AJCC) ^18^. Available clinical data includes patient age, sex, ethnicity, hepatitis B or C infection, tumor staging, resection date, first recurrence data and last follow-up date, as well as tumor budding staging as determined by Budau et al ^19^.

### 2.3 Sample Preparation for LC-MS/MS

All tissue slides were deparaffinized through consecutive baths in xylol, 99 %, 96 %, 70 %, and 50 % ethanol. The slides were stored in water until the tumor and TANM tissue were macrodissected by a pathologist, separately transferred into reaction tubes, and stored at –80°C. For protein extraction, samples were suspended in 0.1 % RapiGest (Waters) and 100 mM 4– (2-hydroxyethyl)-1-piperazineethanesulfonic acid (HEPES), sonicated with a BioRuptor (Diagenode) in 20 cycles à 30 s and heated at 90 °C for 1.5 hours ^34^. Resulting protein concentrations were quantified via a copper / bicinchoninic acid assay (BCA, ThermoFisher Scientific) and up to 100 µg of protein per sample were reduced with 5 mM dithiothreitol (Sigma Aldrich) and incubated in the dark for 30 min. After subsequent alkylation with 10 mM iodoacetamide (Sigma Aldrich) and incubation at 37 °C for 30 min, samples were digested with 1:50 (m/m) lysyl endopeptidase C (Fujifilm) at 42 °C for 2 hours and 1:50 (m/m) trypsin (Worthington) at 37 °C overnight. RapiGest was hydrolyzed by the addition of 2 % trifluoroacetic acid, incubation at 37 °C for 30 min, and 15 min centrifugation at 21,000 x g. The peptide containing supernatant was desalted using a solid phase peptide extraction kit (PreOmics) according to the manufacturer’s instructions. Peptide concentrations were again determined via BCA and 2 µg of peptide per sample were aliquoted for LC-MS/MS measurement and vacuum dried after the addition of 200 fmol of indexed retention time standards (iRTs, Biognosys). In parallel to peptide aliquoting, 1 µg peptides from 24 samples representative for patient sex, age, tumor staging, and TTR distribution were additionally pooled as a reference peptide library.

### 2.4 Patient-derived Xenografts

FFPE tumor sections from nine different patient-derived ICC xenograft (PDX) mouse models were provided by Charles River Germany (am Flughafen 12-14, Freiburg). Tumor tissue was propagated as described before^35^. The study was carried out in accordance with the recommendations of GV-SOLAS in an AAALAC-accredited animal facility. All animal experiments were approved by the Committee on the Ethics of Animal Experiments of the regional council (Permit Numbers I-19/02). ICC PDX models were implanted subcutaneously in 4–6-week-old female NMRI nu/nu mice (Charles River, Sulzfeld) under isoflurane anesthesia. Tumor growth was determined by a two-dimensional measurement with calipers twice a week. Analysis of the PDX models were performed when tumor size reached 800 – 100 mm³. Tumors were sampled, fixed for 24 h in Formalin, and subsequently embedded in paraffin. Deparaffinization was performed as described above. 4 % SDS in 0,1 M HEPES was added to the tissue, which was then sonicated with a BioRuptor in 20 cycles à 40 s and boiled at 95 °C for 90 minutes. Reduction and alkylation were performed by adding 5 mM Tris(2-carboxyethyl)phosphin-hydrochloride and 10 mM 2-choloracetamide with subsequent incubation for 30 min at 37 °C in the dark. After centrifugation, 12 % aqueous phosphoric acid was added to the sample supernatant in a 1:10 ratio. Upon addition of a 3-fold excess volume of 90 % methanol containing 0,1 M triethylammonium bicarbonate (TEAB), the sample was loaded onto an S-Trap micro column (ProTifi) and then washed three times with the same buffer. Trypsin was added in a ratio of 1:10 relative to a sample’s total protein content in a 50 mM TEAB buffer and incubated for 2 h at 47°C. Afterwards, digested peptides were eluted in three steps: first in 50 mM TEAB, second in 0,2% formic acid, and third in 50 % acetonitrile containing 0,2 % formic acid. Subsequently, samples were aliquoted as described above and a library was generated with input from all PDX samples.

### 2.5 LC-MS/MS Measurement

Dried peptides were dissolved in 10 µl aqueous buffer containing 0.3 % acetic acid, sonicated for 5 min, and centrifuged for 5 min at 21,000 x g. The supernatant was transferred into vials and 5 µl, equivalent to 1 µg of peptides, were injected into an Easynlc 1000 nano-flow HPLC coupled to a Q-Exactive Plus mass spectrometer (both ThermoFisher Scientific). Chromatographic separation was performed via a 120 min gradient at 300 nl/min with aqueous buffers A containing 0.3 % acetic acid and B containing 80 % acetonitrile with 0,3 % acetic acid. A self-packed 30 cm pico-frit column (New Objective) with an inner diameter of 100 µm, containing Reprosil Pur 3 µm C18 beads with 100 Å pore size (Dr. Maisch) was used. The gradient was running from 5 to 8 % buffer B for 3 min, increased to 43 % B within 90 min, then to 65 % B within 20 min, and to 100 % B in 5 min, and ended with 2 more minutes washing at 100 % B. Both peptide library and sample measurements were performed in data independent acquisition (DIA) mode^36,37^. Peptide library acquisition was performed via six gas-phase fractionation measurements, each covering a 100 m/z range between 400 and 1000 m/z, consisting of 25 staggered acquisition windows of 4 m/z. Samples were measured in a scan range from 385 to 1015 m/z in staggered acquisition windows of 24 m/z each. All measurements were performed with a resolution of 17,500 and a maximum injection time of 80 ms. Peptides were fragmented via Higher-energy C-trap dissociation (HCD) at stepped normalized collision energies of 25 and 30.

PDX samples were measured with the same setup and settings. However, peptides were separated on a μPAC 2m reverse-phase C18 column (ThermoFisher Scientific, USA) via a gradient going from 8 to 10 % buffer B in 2 min, then to 20 % B within 22 min, to 40 % B in 46 min, to 55 % B in 10 min, and finally to 100 % B in 2 min. There it remained for another 38 min.

### 2.6 Data Analysis

Library generation and peptide-to-spectrum matching were performed in DIA-NN version 1.7 ^38^ by using a human proteome datafile containing reviewed sequences as downloaded from Uniprot on September 1^st^, 2020. Protein quantification was performed via the MaxQuant label-free quantification (LFQ) algorithm as implemented in DIA-NN. For library generation, files were processed with 10 % false discovery rate (FDR), while spectral reannotation was performed with 1 % FDR. In both cases, up to 1 missed cleavage site was allowed and peptides ranging from 7 to 30 amino acids length were included. To analyze proteolytic activity, library files were refined against a human proteome database containing all possible semi-specific peptide sequences. 1 % FDR was applied and no missed cleavage sites were permitted for library generation and reannotation of peptides of 7 to 30 amino acids length.

All subsequent data analysis was performed in R using RStudio and in-house scripts. The DIA-NN output was exported as an expression matrix by using the DIA-NN package ^39^ and then log2 transformed, median-normalized, and corrected for measurement batch effects via ComBat^40^. Hierarchical clustering and principal component analysis (PCA) were performed with MixOmics, Monte-Carlo simulation with M3C ^41,42^. For linear modeling we used the Limma package^43^. Network analyses were performed using Cytoscape ^44^ and enrichment analyses via ClusterProfiler^45^ using KEGG^46,47^, Reactome ^48^, and GeneOntology ^49,50^ databases. Survival statistics, including Cox proportional hazards model, were applied via the survival ^51^ and survminer ^52^ packages.

Analysis of proteolytic processing was performed using in-house-developed R scripts for the annotation and visualization of semi-specific peptides (publicly available on GitHub as the Fragterminomics package) ^53^.

Xenograft samples were analyzed via DIA-NN, the DIA-NN R package and MixOmics as described above ^38,41^. Here, a combined human-mouse proteome datafile as downloaded from Uniprot on September 1^st^, 2020 was used for reannotation.

## 3 Results and Discussion

### 3.1 Description of Cohort

Our ICC cohort comprised 80 tumor tissues and 67 patient-matched, non-malignant adjacent tissues (TANM) from a total of 80 ICC patients. Patient characteristics are summarized in Table 1. The size of our ICC cohort is smaller than the Fudan University ICC (FU-ICC) cohort, for which comprehensive proteogenomic characterizations have recently been published by Dong et al and Lin et al ^26,27^. However, disease etiology and incidence are assumed to differ across world regions and to our knowledge, this study represents the first large proteomic analysis of a western cohort^1,54^.

**Table 1:**
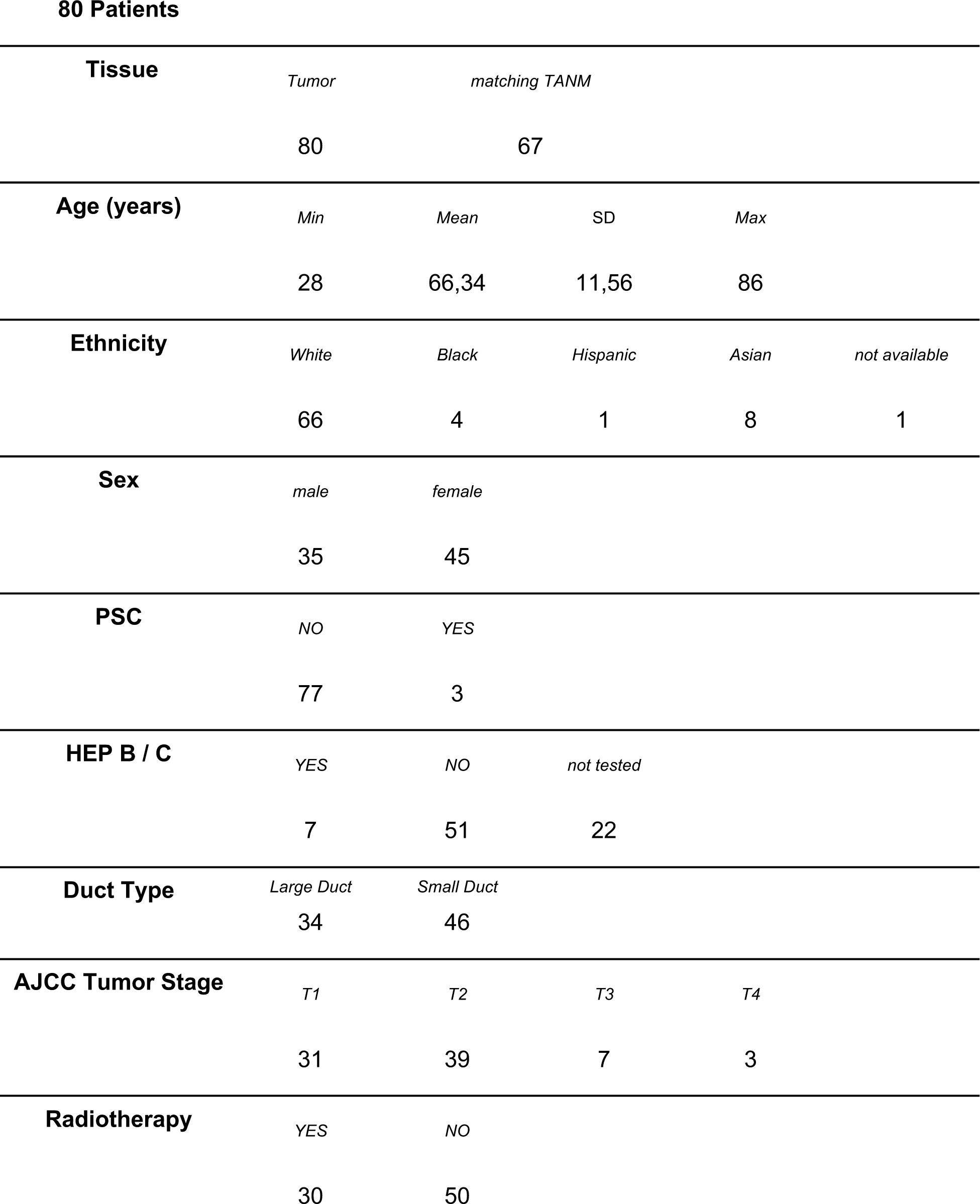
Clinical parameters. PSC – Primary Sclerosing Cholangitis. HEP B/C – diagnosed Hepatitis B or C infection.

### 3.2 Proteome Coverage

Across the cohort, on average 2,330 proteins were identified per sample through DIA measurements on a Thermo QExactive Plus (Fig. 1B,C). We have used DIA-NN with GPF-refinement for data analysis as our recent benchmarking study has underlined the adequacy of this approach ^32^. More than 1,700 proteins presented with less than 20% missing values throughout the entire cohort and were used for statistical analysis (Fig. 1D). To account for putative measurement batch effects, we employed the ComBat ^40^ algorithm for correction, which led to a clearer separation of tumor and TANM tissue in principal component analysis (PCA) (Fig. 1E, F). Six outliers in this analysis were reevaluated by a pathologist, reclassified as hepatocellular carcinomas or fibrosis, and removed from the study; yielding the aforementioned numbers of 80 ICC tumors and 67 TANM tissues. In a hierarchical clustering of the entire cohort, tumor and TANM formed two separate subgroups. Neither digestion and measurement batches, nor the measurement order correlated with the clustering result or the TTR distribution, which indicates successful sample randomization and batch correction (Fig. 1G, S1). The proteome coverage of our study remains below recently reported proteogenomic profiles ^26,27^. We explain these differences with the usage of FFPE specimens and the omission of peptide fractionation.

### 3.3 Tumor and TANM Tissues Show Highly Different Proteome Profiles

Principal component analysis of ICC and TANM proteomes suggests highly divergent proteome biology (Fig. 1F). In order to determine differentially regulated proteins in ICC vs. TANM, we have employed linear models as implemented in the Limma R package ^43^. This approach revealed 1,368 proteins to be differentially regulated (p < 0.01, with FDR correction) (Fig. 2A). We sought to identify common biological themes within this group of proteins. To this end, we applied KEGG enrichment analysis. Proteins connected to Fc-γ-mediated phagocytosis, as well as translation, splicing, and intracellular transport machineries were upregulated in ICC (Fig 2B, S2). In a network analysis, an array of splicing– and DNA-repair associated proteins (including Poly-[ADP-ribose]-polymerase 1 – PARP1) coordinated around Oncogene FUS also showed increased expression within ICCs (Fig. 2C). Enrichment of Fc-γ-mediated phagocytosis, spliceosome, and DNA-Repair with concurrent down-regulation of proteasome components was previously described by Dong et al. as a trans-effect of a characteristic chromosome 14q loss ^26^. We also identified extracellular matrix (ECM) and matrisome components (e.g. Biglycan, Lumican, and various collagens) to be among the most prominently increased proteins within ICC tumor tissue, which is in agreement with strong desmoplastic reactions being a hallmark of ICC ^22^. Compared to TANM tissue, ICC proteins belonging to processes such as citrate cycle, oxidative phosphorylation, and proteasomal degradation were downregulated (Fig 2B, S2). In part, this observation adheres to the commonly reported Warburg effect within solid tumors ^55^.

**Figure 2.**
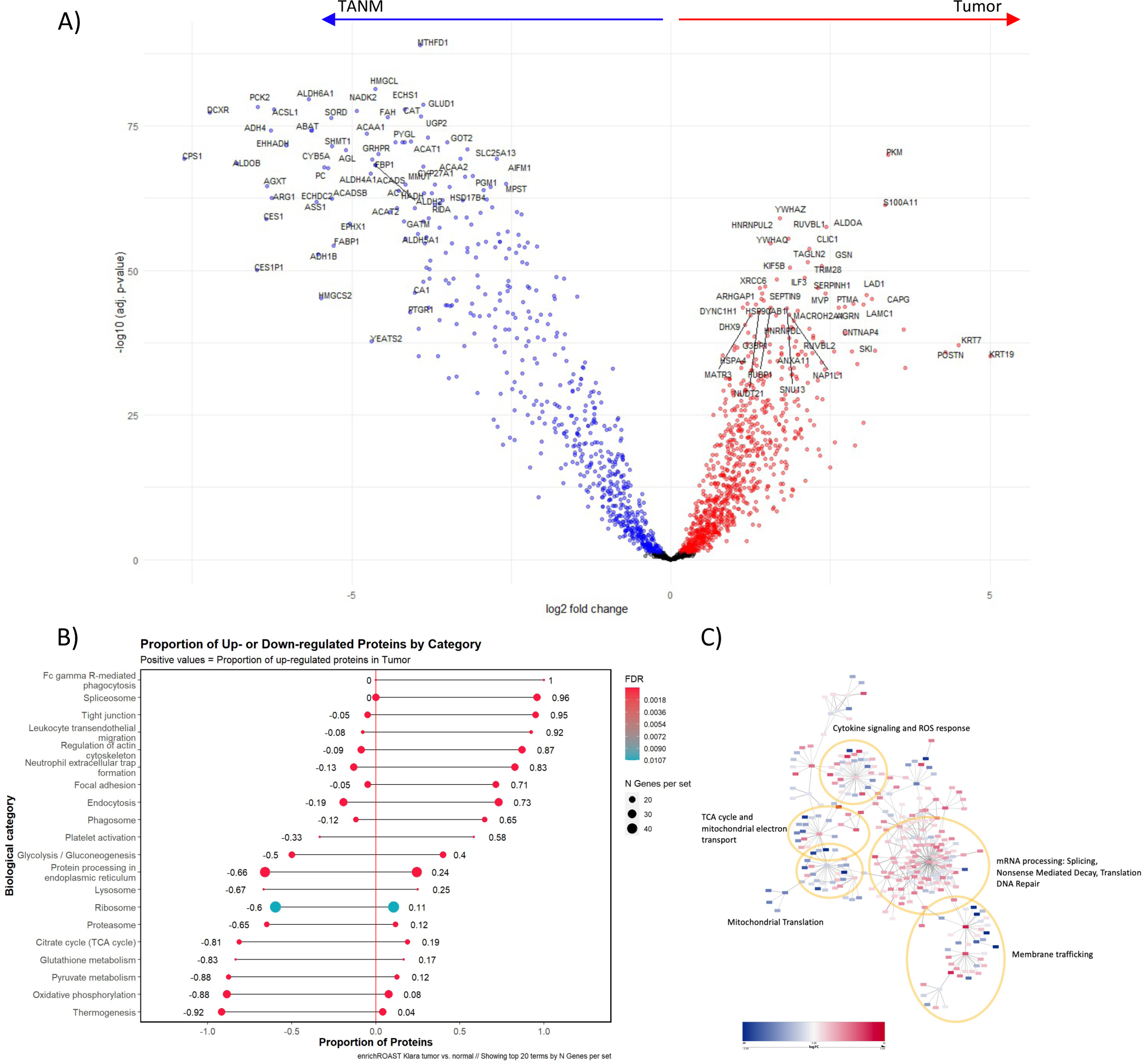
– tumor vs. TANM tissue: A) Volcano plot of differentially expressed proteins between tumor and TANM tissue. B) KEGG enrichment analysis of differentially regulated biological processes between tumor and TANM. C) Protein interaction network of differential expression analysis results. Log fold changes > 0 in red indicate high expression in tumor, log fold changes < 0 in blue high expression in TANM.

### 3.4 ICC Tumor Proteomes form Two Clusters with Strong Proteome Differences and Diverging Times-to-Recurrence

Recent proteome studies on solid tumors ^28–30^ including ICC ^26^, have highlighted the existence of proteomic subgroups within a tumor entity; i.e. subgroups that are enriched in proteins stemming from (tumor-) biological processes. Unsupervised PCA of our ICC proteome data showed homogenous distribution along components 1 and 2, and did not follow trends regarding sex, age, or tumor stage. Next, we performed unsupervised hierarchical clustering, which sorted tumors into two proteomic clusters (Fig. 3A), which was confirmed through a Monte-Carlo simulation. Linear modeling of differentially expressed proteins followed by gene set enrichment via KEGG and REACTOME databases revealed different biological processes to be upregulated in each cluster. In cluster 1, we detected a fingerprint of matrisomal proteins, focal adhesion, and coagulation and complement cascades. Cluster 2 exhibited increased activity in synthesis, processing, and degradation of RNA and proteins (“RNA– and protein turnover”), and glucose metabolism (Fig. 3B&C, S3). We further investigated matrisomal proteins identified in cluster 1 and found elevated expression to be confined to ECM Glycoproteins, Collagens, and Proteoglycans (Fig. 4).

**Figure 3.**
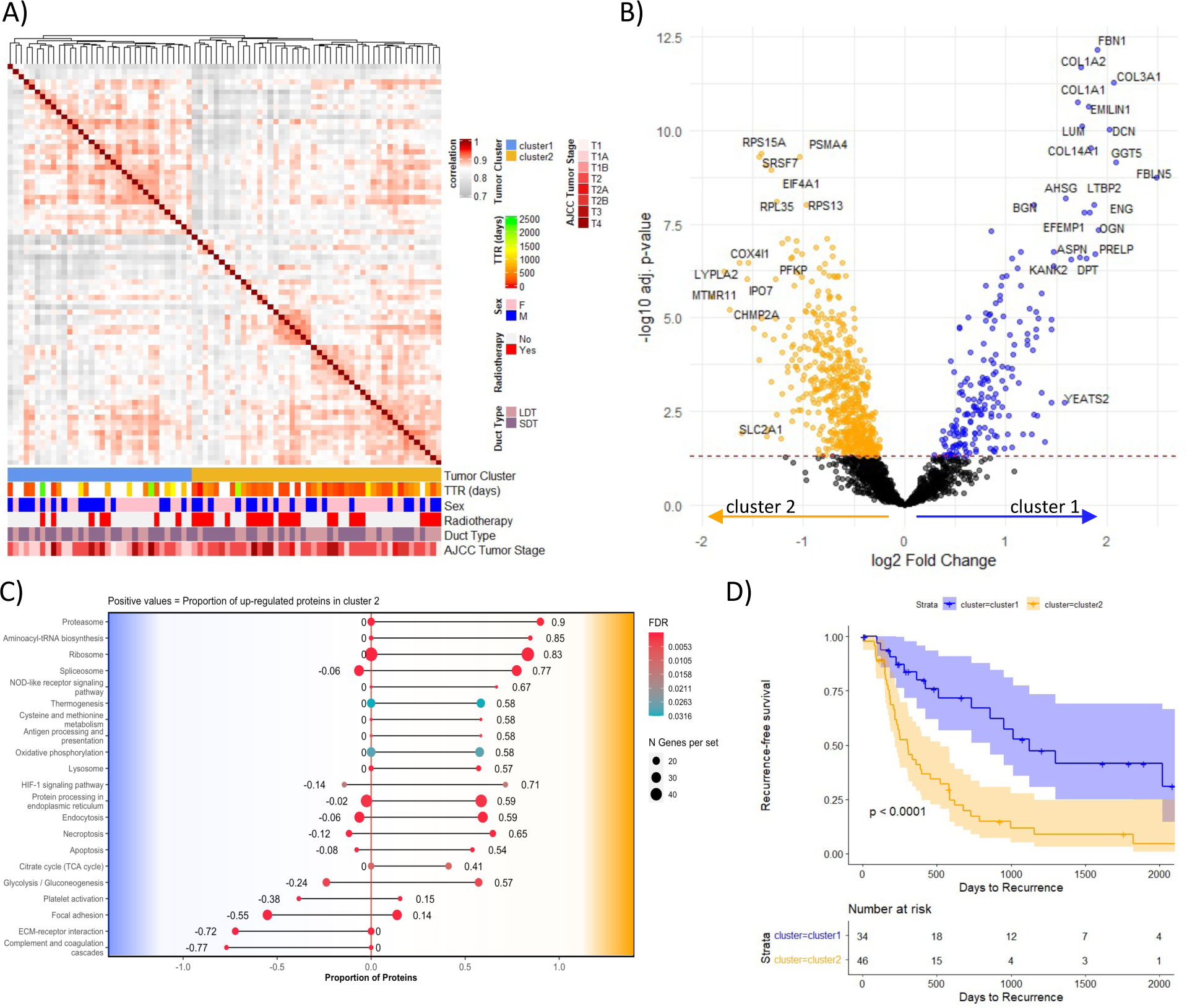
– hierarchical clustering: A) Correlation plot indicating Pearson-correlation of hierarchically clustered tumor samples. Matching clusters, TTR length, sex, radiotherapy treatment, duct type, and AJCC tumor stage are indicated below. B) Volcano plot of differentially expressed proteins. C) KEGG enrichment analysis of differentially regulated biological processes between both clusters. D) Kaplan-Meier curve comparing TTR distribution between cluster 1 and 2.

**Figure 4.**
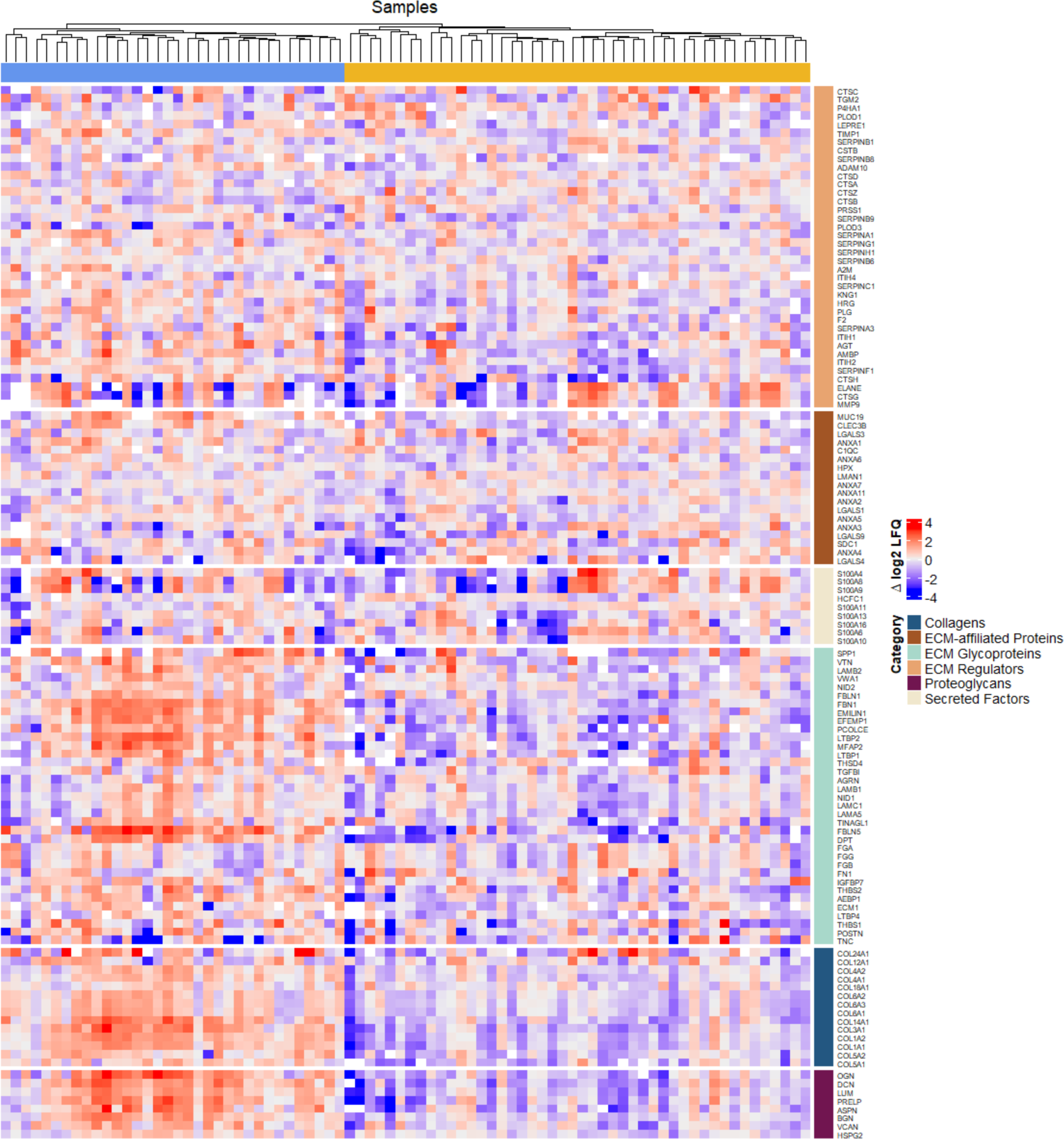
– ECM protein expression: Heatmap of ECM protein expression in hierarchically clustered tumor samples.

Mosele et al. have compiled a list of highly relevant, clinically actionable target genes/proteins (HATs) in the context of solid tumors ^56^. We have assessed whether the two subclusters present a differential abundance of HATs. However, only cytoplasmic and mitochondrial isocitrate dehydrogenase 1 and 2 (IDH1 & IDH2) were found in our dataset, but without differential abundance between the two subgroups.

Next, we probed whether both clusters differ in clinicopathological features. Most clinical parameters listed in Table 1, including sex and duct type classification, were equally distributed across both clusters (Fig. 3A, Table 2). However, cluster 2 presented with a slightly higher fraction of AJCC stage T2 specimens, while cluster 1 included more T1 tumors. Most importantly, cluster 1, enriched with ECM proteins, is associated with significantly prolonged TTR, while cluster 2, enriched with protein turnover components, is associated with a significantly shortened TTR (Fig. 3D).

**Table 2:** Clinical annotation of subclusters. PSC – Primary Sclerosing Cholangitis. HEP B/C – diagnosed Hepatitis B or C infection.

|  | cluster 1 |  |  |  |  | cluster 2 |  |  |  |  |
| --- | --- | --- | --- | --- | --- | --- | --- | --- | --- | --- |
| 80 patients | 34 |  |  |  |  | 46 |  |  |  |  |
| Age (years) | Min | Mean | SD | Max |  | Min | Mean | SD | Max |  |
|  | 40 | 66,06 | 9,62 | 81 |  | 28 | 66,54 | 12,9 | 86 |  |
| Ethnicity | White | Black | Hispanic | Asian | not available | White | Black | Hispanic | Asian | not available |
|  | 28 | 2 | - | 3 | 1 | 38 | 2 | 1 | 5 | - |
| Sex | male | female |  |  |  | male | female |  |  |  |
|  | 16 | 18 |  |  |  | 19 | 27 |  |  |  |
| PSC | NO | YES |  |  |  | NO | YES |  |  |  |
|  | 33 | 1 |  |  |  | 44 | 2 |  |  |  |
| HEP B / C | YES | NO | not tested |  |  | YES | NO | not tested |  |  |
|  | 5 | 20 | 9 |  |  | 2 | 31 | 13 |  |  |
| Duct Type | Large Duct |  | Small Duct |  |  | Large Duct |  | Small Duct |  |  |
|  | 13 |  | 21 |  |  | 21 |  | 25 |  |  |
| Tumor Stage | T1 | T2 | T3 | T4 |  | T1 | T2 | T3 | T4 |  |
|  | 18 | 11 | 3 | 2 |  | 13 | 28 | 4 | 1 |  |
| Radio-therapy | YES | NO |  |  |  | YES | NO |  |  |  |
|  | 7 | 27 |  |  |  | 23 | 23 |  |  |  |

These results are largely in line with previous multi-omics publications. Dong et al describe four ICC subclusters based on mRNA, proteomic– and phosphoproteomic data for the FU-ICC cohort ^26^. FU-ICC cluster 1 was abundant with proteins involved in inflammation and glucose metabolism. Conversely, FU-ICC cluster S2 showed high levels of ECM proteins. FU-ICC cluster 3 was enriched with proteins involved in MAP-Kinase signaling and fatty acid and nucleotide synthesis, while FU-ICC cluster 4 biology remained inconclusive. FU-ICC cluster 1 had the worst prognosis, particularly for tumors of AJCC stages I and II. Similarly, Lin et al report three immune subgroups within the same cohort and link immune-activated cluster IG3 (fibroblast and lymphocyte invasion) to prolonged survival, while other clusters IG1 and IG2 are characterized by a worse prognosis and innate immunity mediated inflammation (IG1) or immune exclusion (IG2) ^27^.

### 3.5 TANM Proteome Clusters are Independent from ICC Clusters

Similar to the tumor samples, we sought to identify distinct proteome clusters of TANM tissue. Hierarchical clustering yielded three distinct clusters (TANM-cluster) as further corroborated by Monte-Carlo simulation. Linear modeling followed by enrichment analysis revealed TANM-cluster 1 to express proteins for glucose metabolism and oxidation, TANM-cluster 2 for complement activation, and the small TANM-cluster 3 for coagulation, perhaps as a result of bleeding during surgery. The TANM-clusters do not correlate with the TTR, sex, duct type, tumor stage, or radiotherapy, and do not match to the clustering of corresponding ICC tumors (Fig. 5, S4).

**Figure 5.**
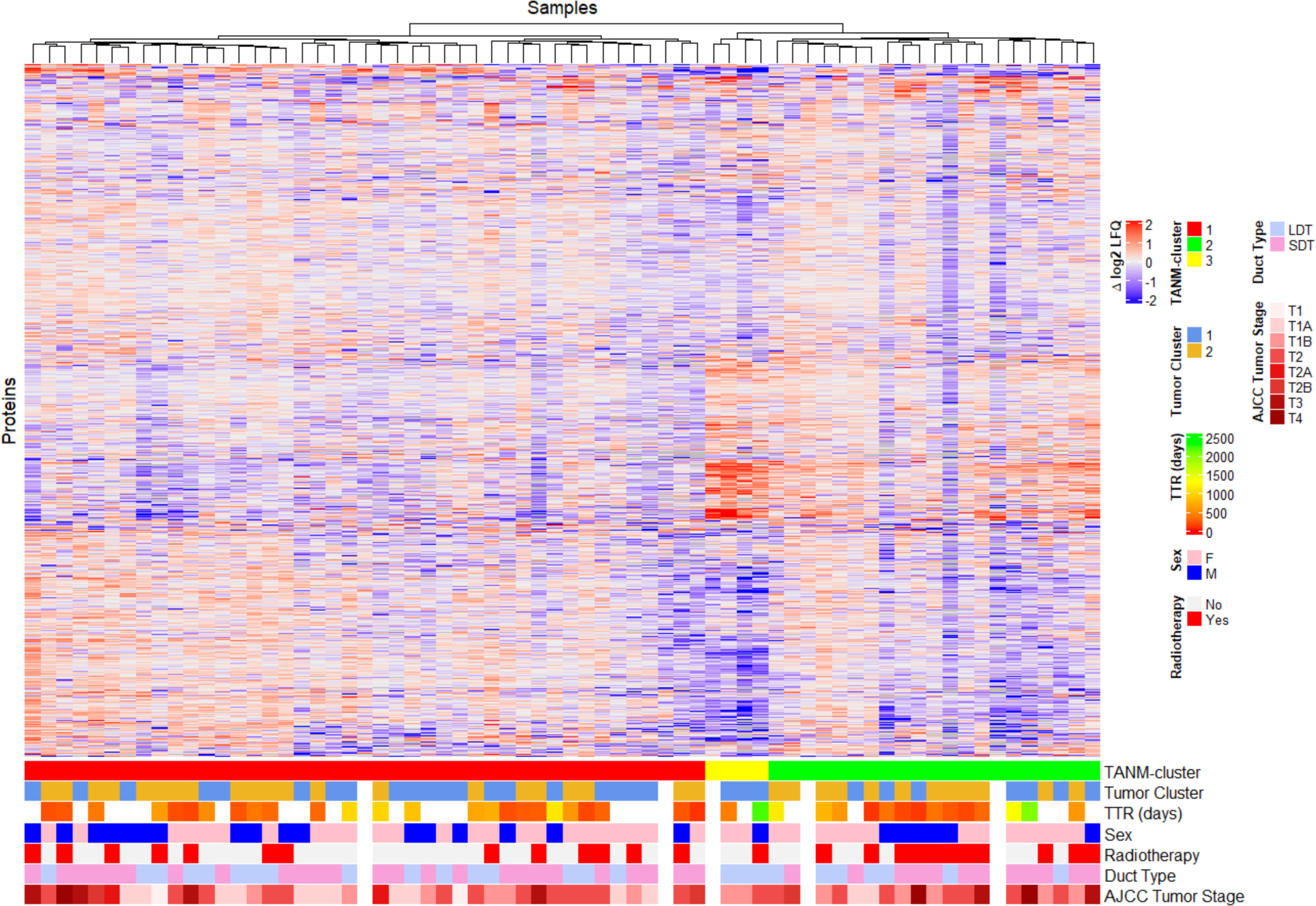
– TANM tissue: Heatmap of hierarchically clustered TANM samples. Matching TANM cluster, tumor cluster, TTR, sex, radiotherapy, duct type, and AJCC 8th Edition tumor stage are indicated below.

### 3.6 Cox Proportional Hazards Model Highlights Prognostic ICC Proteins

In a second analysis, we iteratively applied Cox proportional hazards model (CPHM) to each protein by including adjuvant radiotherapy as covariable. Radiotherapy was significantly correlated with a worse TTR prognosis (Fig. 6A,B). After correction for multiple testing, we identified individual proteins whose abundance correlates with TTR distribution. Samples of patients who did not experience a recurrence throughout the study were censored and included in the analysis. The resulting list of protein candidates was adjusted for multiple-testing errors via Benjamini-Hochberg false discovery rate correction. 39 proteins were found to have expression patterns significantly correlating to the TTR distribution (Fig. 6C). Of these, 12 showed a hazard ratio above 1, indicating an increased risk for early recurrence. Gene set enrichment analysis via the KEGG database revealed a large fraction of proteins with high hazard ratios to be involved in cell-cycle progression and protein turnover (Fig. 6D). Conversely, 27 proteins were associated with hazard ratios below 1 and a decreased risk for early recurrence. Many of the proteins with hazard ratios below 1 appeared to be involved in ECM processes like focal adhesion (Fig. 6E). A majority of the 39 detected proteins also showed significant expression changes over the TTR, with particularly pronounced differences in Tensin 1 (TNS-1), Inter-alpha-trypsin inhibitor heavy chain H2 (ITIH2), Histone 1.0 (H1-0), Biglycan (BGN), Vitronectin (VTN), Lumican (LUM), Glutathione Hydrolase 5 (GGT5), Fibrillin-1 (FBN1), Collagen alpha-1(III) chain (COL3A1), Prolargin (PRELP), L-lactate dehydrogenase A chain (LDHA), SKI oncoprotein (SKI), and Proteasome subunit beta type-4 (PSMB4 – all Table 3). 31 of these proteins, including the ones mentioned, were also significantly altered between cluster 1 and 2. Moreover, in this independent analysis, similar biological processes were associated with high or low risk for recurrence as in the previous hierarchical clustering. Interestingly, patients’ overall survival did not correlate well with protein expression in the CPHM, although both clusters also are significant predictors (Fig. S5). This is in line with Hyder et al. who noted that prognostic factors for the time to recurrence do not necessarily correlate with overall survival ^57^.

**Figure 6.**
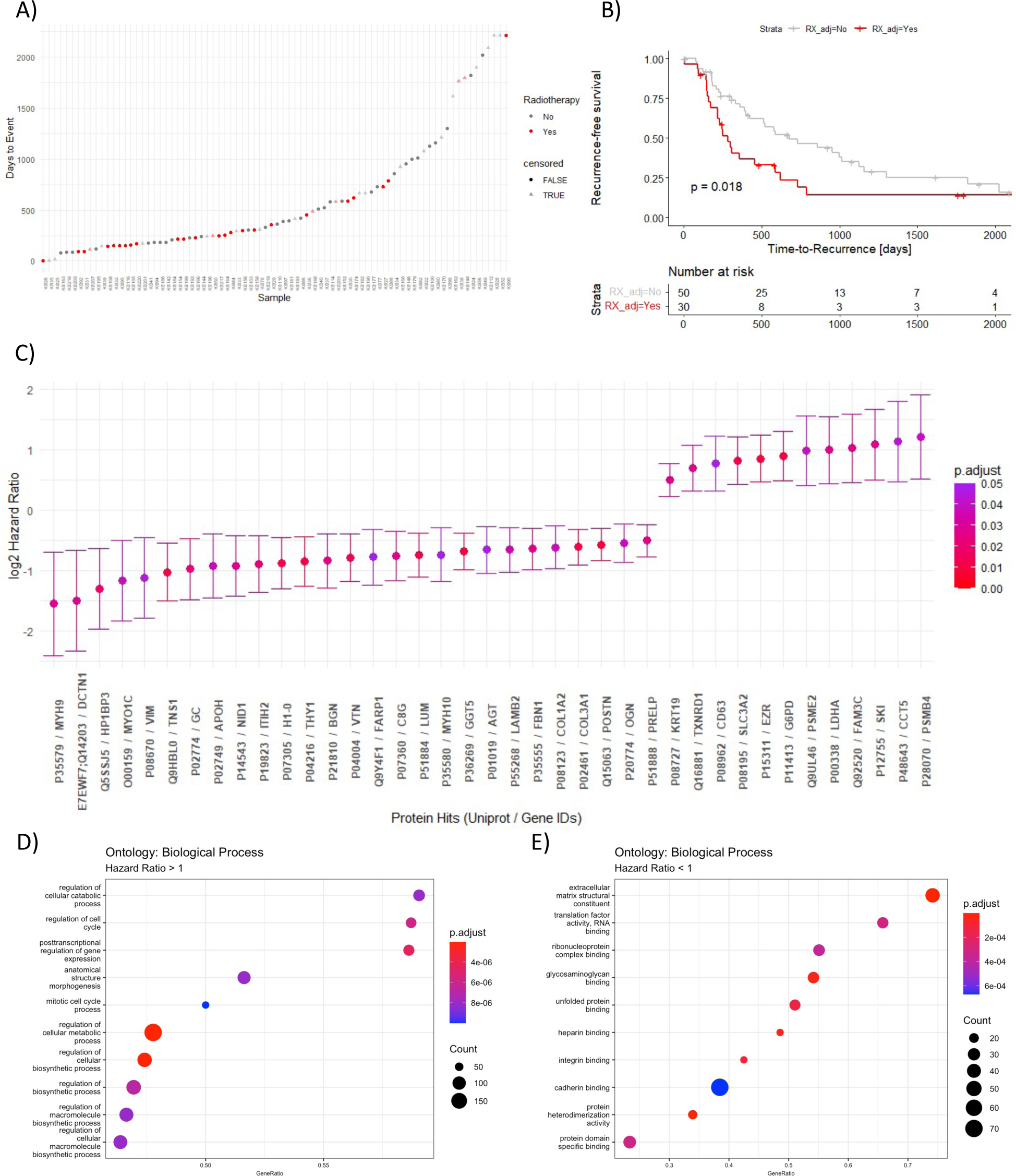
– survival statistics: A) TTR distribution across the cohort. B) Kaplan-Meier curve comparing patients with radiotherapy (RX_adj) to patients without. C) CPHM significant protein hits with radiotherapy as covariable. D, E) KEGG enrichment analyses of upregulated biological processes among proteins with high and low hazard ratios.

**Table 3:** CPHM result.

| **UNIPROT ID** | **Hazard Ratio** | **p-value** | **Adjusted p-value** | **Gene** | **Protein Name** |
| --- | --- | --- | --- | --- | --- |
| **P35579** | 0.34 | 0.00039 | 0.02788 | MYH9 | Myosin-9 |
| **Q14203** | 0.35 | 0.00038 | 0.02788 | DCTN1 | Dynactin subunit 1 |
| **Q5SSJ5** | 0.40 | 0.00012 | 0.01809 | HP1BP3 | Heterochromatin protein 1-binding protein 3 |
| **O00159** | 0.45 | 0.00062 | 0.03818 | MYO1C | Unconventional myosin-Ic |
| **P08670** | 0.46 | 0.00087 | 0.04516 | VIM | Vimentin |
| **Q9HBL0** | 0.49 | 0.00003 | 0.01029 | TNS1 | Tensin-1 |
| **P02774** | 0.51 | 0.00017 | 0.02257 | GC | Vitamin D-binding protein |
| **P02749** | 0.52 | 0.00063 | 0.03818 | APOH | Beta-2-glycoprotein 1 |
| **P14543** | 0.53 | 0.00028 | 0.02788 | NID1 | Nidogen-1 |
| **P19823** | 0.54 | 0.00016 | 0.02257 | ITIH2 | Inter-alpha-trypsin inhibitor heavy chain H2 |
| **P07305** | 0.54 | 0.00006 | 0.01152 | H1-0 | Histone H1.0 |
| **P04216** | 0.56 | 0.00004 | 0.01152 | THY1 | Thy-1 membrane glycoprotein |
| **P21810** | 0.56 | 0.00023 | 0.02566 | BGN | Biglycan |
| **P04004** | 0.58 | 0.00009 | 0.01470 | VTN | Vitronectin |
| **Q9Y4F1** | 0.58 | 0.00103 | 0.04872 | FARP1 | FERM, ARHGEF and pleckstrin domain-containing protein 1 |
| **P07360** | 0.59 | 0.00031 | 0.02788 | C8G | Complement component C8 gamma chain |
| **P51884** | 0.60 | 0.00006 | 0.01152 | LUM | Lumican |
| **P35580** | 0.60 | 0.00104 | 0.04872 | MYH10 | Myosin-10 |
| **P36269** | 0.62 | 0.00001 | 0.00963 | GGT5 | Glutathione hydrolase 5 proenzyme |
| **P01019** | 0.63 | 0.00097 | 0.04796 | AGT | Angiotensinogen |
| **P55268** | 0.63 | 0.00052 | 0.03532 | LAMB2 | Laminin subunit beta-2 |
| **P35555** | 0.64 | 0.00024 | 0.02599 | FBN1 | Fibrillin-1 |
| **P08123** | 0.65 | 0.00061 | 0.03818 | COL1A2 | Collagen alpha-2(I) chain |
| **P02461** | 0.65 | 0.00006 | 0.01152 | COL3A1 | Collagen alpha-1(III) chain |
| **Q15063** | 0.67 | 0.00002 | 0.00963 | POSTN | Periostin |
| **P20774** | 0.68 | 0.00065 | 0.03818 | OGN | Mimecan |
| **P51888** | 0.70 | 0.00022 | 0.02566 | PRELP | Prolargin |
| **P08727** | 1.41 | 0.00036 | 0.02788 | KRT19 | Keratin, type I cytoskeletal 19 |
| **Q16881** | 1.62 | 0.00036 | 0.02788 | TXNRD1 | Thioredoxin reductase 1, cytoplasmic |
| **P08962** | 1.71 | 0.00097 | 0.04796 | CD63 | CD63 antigen |
| **P08195** | 1.77 | 0.00005 | 0.01152 | SLC3A2 | 4F2 cell-surface antigen heavy chain |
| **P15311** | 1.80 | 0.00002 | 0.00963 | EZR | Ezrin |
| **P11413** | 1.85 | 0.00002 | 0.00963 | G6PD | Glucose-6-phosphate 1-dehydrogenase |
| **Q9UL46** | 1.98 | 0.00072 | 0.03993 | PSME2 | Proteasome activator complex subunit 2 |
| **P00338** | 1.99 | 0.00038 | 0.02788 | LDHA | L-lactate dehydrogenase A chain |
| **Q92520** | 2.03 | 0.00040 | 0.02788 | FAM3C | Protein FAM3C |
| **P12755** | 2.12 | 0.00033 | 0.02788 | SKI | Ski oncogene |
| **P48643** | 2.20 | 0.00082 | 0.04393 | CCT5 | T-complex protein 1 subunit epsilon |
| **P28070** | 2.31 | 0.00072 | 0.03993 | PSMB4 | Proteasome subunit beta type-4 |

Several of the identified prognostic proteins have been previously associated with the tumor biology of cholangiocarcinoma. For example, high LDHA expression is associated with a poor prognosis, while FAM3C expression was linked to highly malignant ICC cancer stem-cells ^58,59^. Considering SKI oncoprotein, two transcriptomic studies identified its role in the development of ICC ^60,61^.

### 3.7 Endogenous Proteolytic Processing is increased in cluster 1

Increased proteolysis is a property of many cancers, as tumor progression is also facilitated by the decomposition of extracellular structures. Endogenous proteolysis yields truncated peptides which are amenable to proteomic analysis ^62^. We have recently shown that so-called semi-specific approaches in peptide-to-spectrum matching are a valuable approach to grasping endogenous proteolytic processing ^63–65^. Since all samples were digested with trypsin, which specifically cleaves protein sequences after R or K, any detected peptide sequence that deviated from this specificity might represent a physiological cleavage product. Across the cohort, 5.1% (868) of all identified peptides possessed such semi-tryptic properties (Fig. 7A). Interestingly, while we expected to find higher portions of semi-specific peptides in a majority of tumor samples, only tumors associated with ECM-enriched cluster 1 showed significantly increased ratios of semi-specific peptides when compared to matching TANM tissue (Fig. 7B). Moreover, proteolytically degraded proteins seem to differ between tumor and TANM tissue, and also among both clusters (Fig. 7C). Recently, it has been demonstrated that antibodies may specifically recognize proteolytically generated neo-epitopes in solid tumors ^66^. Identification of tumor-specific patterns of endogenous proteolytic processing may supplement such efforts.

**Figure 7.**
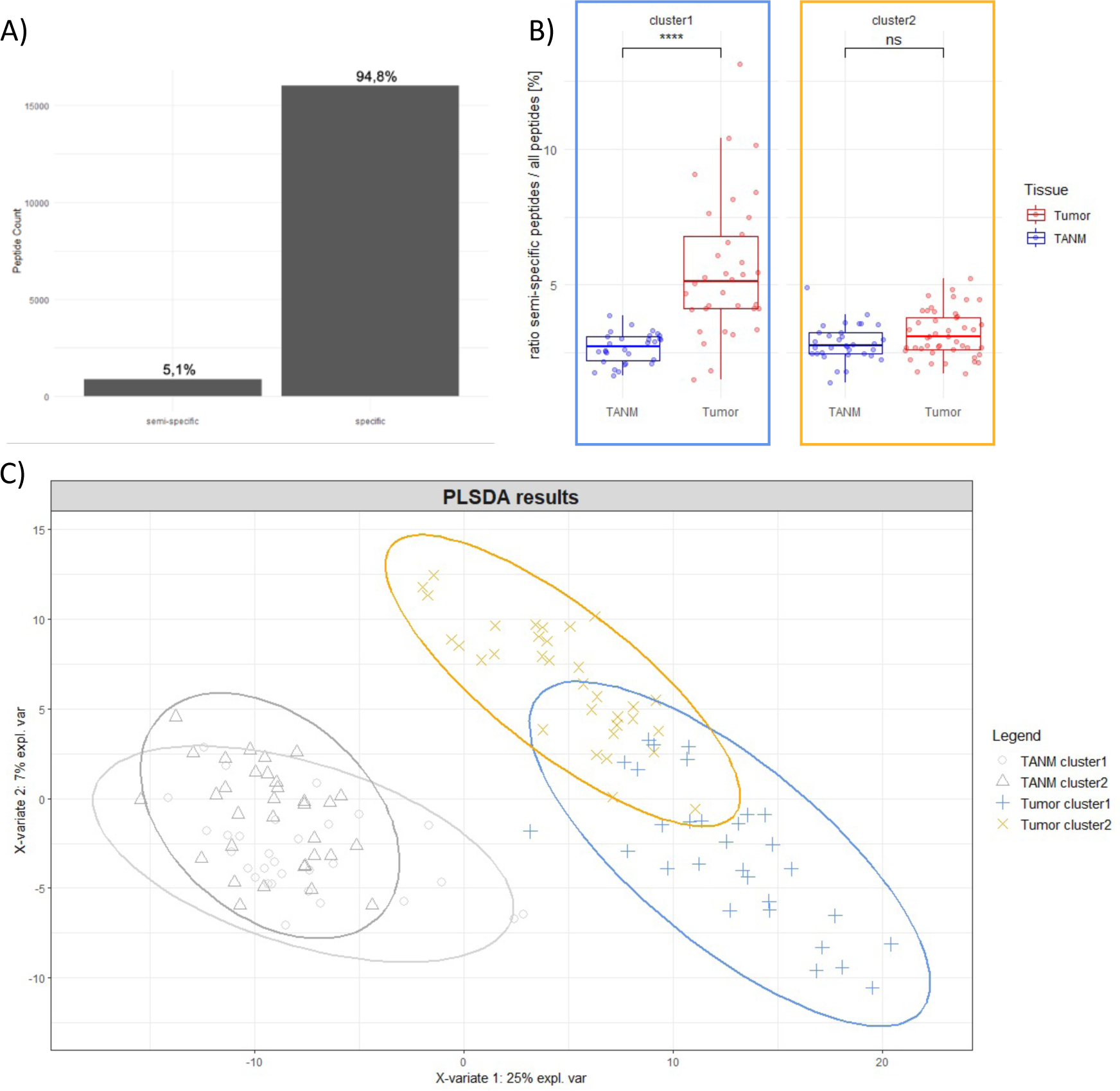
– semi-tryptic analysis: A) Count and percentage of fully tryptic (specific) and semi-specific detected peptides. B) Boxplot indicating the ratio of semi-specific peptides among all detected peptides for tumor and TANM samples in cluster 1 or 2. C) PLS-DA of semi-specific peptide expression in both clusters and matching TANM tissues. Ovals indicate 95% confidence intervals.

### 3.8 Proteomics of ICC PDX Models Enables Insight into Tumor-Stroma Co-Regulation

Given that amino acid sequences between murine and human proteins are different, the species origin of proteins can be inferred. Therefore, patient-derived xenograft (PDX) mouse models offer unique opportunities for studying tumor-stroma interactions ^67^.

In this study, we investigate the interplay of ICC tumors and their surrounding stroma on the proteome level in a small cohort of nine PDX. Since the transplanted tumor was human, invading stromal cells were supposed to be of murine origin. To unambiguously assign human and murine proteins to our DIA data, we only considered peptides with sequences that are unique to mouse and human proteome. On average, we detected 1,491 murine (stroma) and 3,168 human (tumor) proteins per sample (Fig. 8A). These numbers underscore the feasibility of performing mixed-species proteomics experiments with an acceptable proteome coverage. Regarding the interpretation of results, it needs to be considered that xenograft-carrying mice are deficient in adaptive immunity and thus cannot mount T-cell mediated antitumor responses.

**Figure 8.**
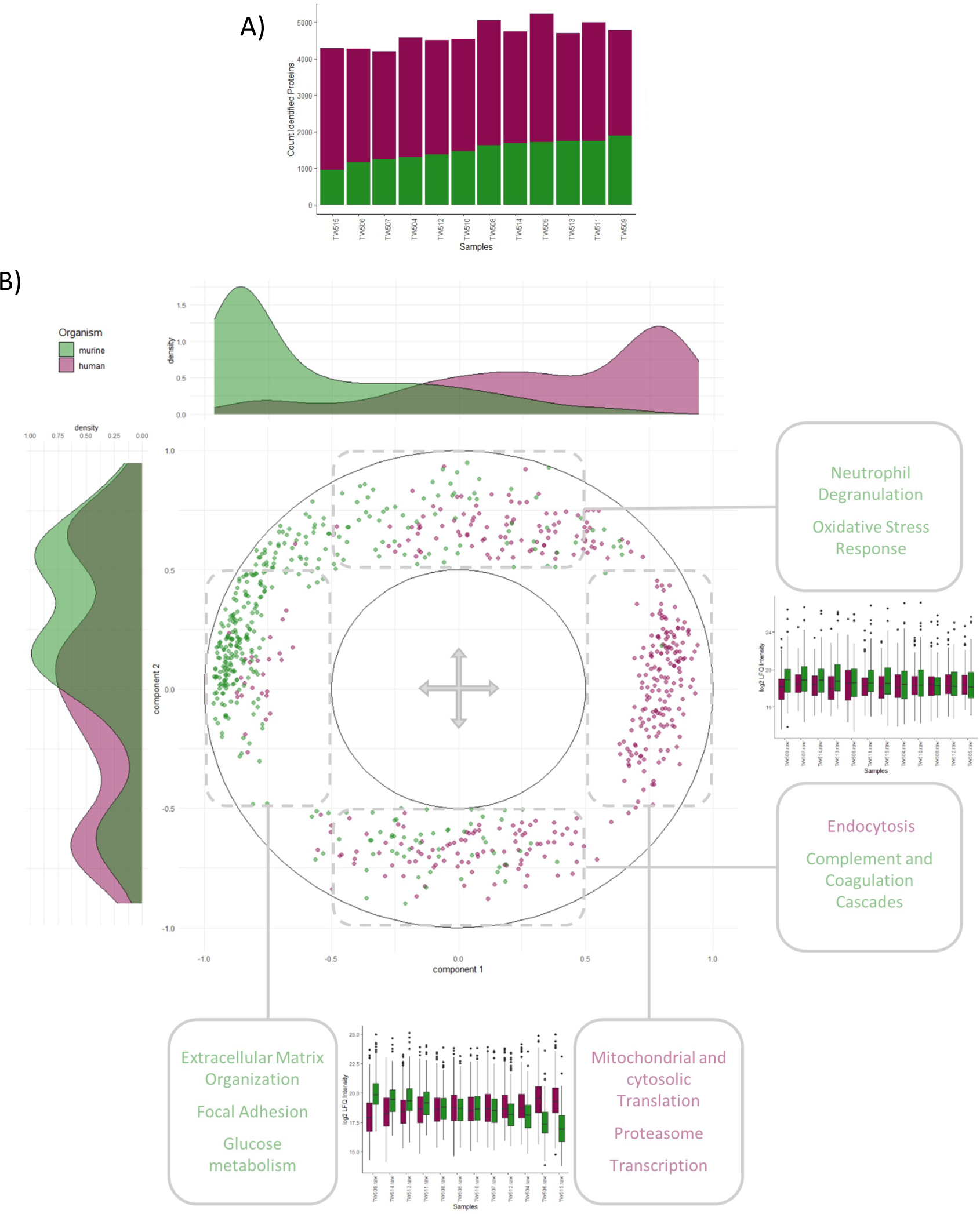
– xenografts: A) Barplot of all detected protein groups per sample. B) Circle plot detailing an unsupervised PLS analysis of the top 200 proteins with the highest factor loadings. Proteins with similar expression patterns cluster together. Proteins with inverse expression patterns aggregate at opposing axis ends. Component 1 represents the strongest contrast in expression regulation, component 2 the second strongest. Reactome enrichment results for proteins clustering at each axis end are provided in rounded boxes. Density plots illustrate the distribution of human and mouse proteins across both components, while boxplots indicate each component’s human vs. mouse protein content per sample.

We generated an unsupervised partial least squares (PLS) model of the data, defining co– or inversely regulated sets of proteins – independently of their species-origin. For the first two components, we focused on the top 200 proteins with the highest absolute factor loadings. By this approach, we could identify proteins sharing similar expression patterns throughout the PDX cohort and functionally compare them to their inversely regulated counterparts. The coregulation is illustrated by a circle plot (Fig. 8B). In brief, proteins aggregating together are showing similar expression patterns, whereas proteins on opposing ends of either axis are inversely regulated.

Unsurprisingly, murine and human proteins each form contrasting sets along component 1, meaning that the strongest inverse expression pattern is determined by species origin. An enrichment analysis of component 1 showed mainly murine proteins associated with neutrophil degranulation, ECM organization, cellular respiration, as well as glucose and amino acid metabolism on the negative side. On the opposing, positive side we found human proteins involved in mitochondrial and cytosolic translation, ribosome biogenesis, and protein metabolism. This divergent expression pattern reflects identified clusters of the human ICC cohort and supports the notion that innate immune reactions and ECM deposition are initiated by stromal cells invading the tumor. Accordingly, we observe a similar contrast in protein expression along component 2. At the positive axis end, human proteins were enriched for oxidative stress response and metabolism (mainly of lipids). At the negative axis end, murine proteins linked to ECM organization and collagen synthesis were increased. Since the mouse models for this PDX cohort were isogenic, genomic diversity was confined to PDX tumor cells. As observed in our cohort and these xenografts, different translational and metabolic activities between tumors might direct stromal reactions. Additionally, murine homologues of many cluster-1-associated proteins in the human cohort aggregated on the negative sides of components 1 and 2.

## 4 Conclusion

Here we describe one of the first proteomic characterizations of a western ICC cohort. Matching tumor and TANM tissues from 80 patients showed strong differences, particularly in the expression of splicing, translation, intracellular transport, and ECM proteins, which were increased in tumor tissue. Within the tumor samples, we identified two subgroups with significantly diverging TTR in a hierarchical clustering analysis. Cluster 1, with a beneficial prognosis, was enriched for ECM components, and worse performing cluster 2 for proteins linked to RNA– and protein turnover. In an analysis of non-tryptic protein cleavage events, we also found increased proteolytic activity in tumors of cluster 1, but not in matching TANM tissue and neither in tumors or TANM associated with cluster 2. An independent survival analysis detected similar proteins and biological motifs to correlate with the cohort’s TTR distribution and highlighted the beneficial prognosis for tumors with high ECM content. Interestingly, TANM tissue subclusters did not match to their respective tumor counterparts. Lastly, we investigated tumor-stroma interactions through patient-derived ICC xenografts. Here, we observe opposing regulation of tumoral proteins involved in metabolism and turnover vs. stromal ECM protein; reflecting the major proteomic subtypes of the human ICC cohort.

## 5 Abbreviations

AAALAC: Association for Assessment and Accreditation of Laboratory Animal Care International
AJCC: American Joint Committee on Cancer
BCA: Bicinchonic Acid
BGN: Biglycan
CCA: Cholangiocarcinoma
COL3A1: Collagen alpha-1(III) chain
CPHM: Cox’ Proportional Hazards Model
CPTAC: Clinical Proteomic Tumor Analysis Consortium
DIA: Data Independent Acquisition
DNA: Desoxyribonucleic acid
ECM: Extracellular Matrix
FBN1: Fibrillin-1
FDR: False Discovery Rate
FFPE: Formalin-fixed, Paraffin-embedded
FU-ICC: Fudan University Intrahepatic Cholangiocarcinoma Cohort
FUS: Oncogene FUS
GGT5: Glutathione Hydrolase 5
GV-SOLAS: Gesellschaft für Versuchstierkunde – Society for Laboratory Animal Science
H1-0: Histone 1.0
HAT: Highly Actionable Targets
HCD: Higher-energy Collisional Dissociation
HEPES: 4-(2-hydroxyethyl)-1-piperazineethanesulfonic Acid
HPLC: High-performance Liquid Chromatography
ICC: Intrahepatic Cholangiocarcinoma
IDH1/2: cytoplasmic and mitochondrial isocitrate dehydrogenase 1 and 2
iRTs: Indexed Retention Time Standards
ITIH2: Inter-alpha-trypsin inhibitor heavy chain H2
LC-MS/MS: Liquid-chromatography mass spectrometry
LDHA: L-lactate dehydrogenase A chain
LFQ: Label-free Quantitation
LUM: Lumican
MSKCC: Memorial Sloan Kettering Cancer Center New York
PARP1: Poly-[ADP-ribose]-polymerase 1
PCA: Principal Component Analysis
PDX: Patient-derived Xenograft
PLS: Partial Least Squares
PRELP: Prolargin
PSMB4: Proteasome subunit beta type-4
SDS: Sodiumdodecylsulfate
SKI: SKI oncoprotein
TANM: Tumor-adjacent, non-malignant
TEAB: Tetraethylammonium Bromide
TNS-1: Tensin 1
TTR: Time to Recurrence
UICC: Union for International Cancer Control
VTN: Vitronectin

## 6 Acknowledgments

## 7 Data availability

The proteomic raw data will be available under restricted access for medical data protection reasons upon journal submission. Until then, please contact the corresponding author for access.

## 9 Supplementary Data

– Figure S1: TTR distribution across cohort
– Figure S2: KEGG Tumor vs. TANM
– Figure S3: KEGG enrichment cluster 1 vs. cluster 2
– Figure S4: TANM clusters
– Figure S5: CPHM overall survival
– Figure S6: Xenograft cluster association

## 10 Contributions

OS and PB designed the project; PB, OS, MW, CS, and LT assembled the cohort and collected clinico-pathological data; KB, KK and PB performed pathological assessment; KB and TW processed human samples; TW processed xenograft samples; TW performed mass spectrometric measurements; TW, MCC, FH, and NP analyzed human data; FH, TW, JS, and OS analyzed xenograft data; TW and OS wrote the manuscript.

## List of References

1. Banales, J. M. et al. Cholangiocarcinoma 2020: the next horizon in mechanisms and management. Nat. Rev. Gastroenterol. Hepatol. 17, 557–588 (2020).

2. Banales, J. M. et al. Expert consensus document: Cholangiocarcinoma: current knowledge and future perspectives consensus statement from the European Network for the Study of Cholangiocarcinoma (ENS-CCA). Nat. Rev. Gastroenterol. Hepatol. 13, 261–280 (2016).

3. Rizvi, S., Khan, S. A., Hallemeier, C. L., Kelley, R. K. & Gores, G. J. Cholangiocarcinoma — evolving concepts and therapeutic strategies. Nat. Rev. Clin. Oncol. 15, 95–111 (2018).

4. Khan, S. A., Tavolari, S. & Brandi, G. Cholangiocarcinoma: Epidemiology and risk factors. Liver Int. 39, 19–31 (2019).

5. Clements, O., Eliahoo, J., Kim, J. U., Taylor-Robinson, S. D. & Khan, S. A. Risk factors for intrahepatic and extrahepatic cholangiocarcinoma: A systematic review and meta-analysis. J. Hepatol. 72, 95–103 (2020).

6. Nakanuma, Y. et al. Pathological classification of intrahepatic cholangiocarcinoma based on a new concept. World J. Hepatol. 2, 419–427 (2010).

7. Shin, H.-R. et al. Epidemiology of cholangiocarcinoma: An update focusing on risk factors. Cancer Sci. 101, 579–585 (2010).

8. Kelley, R. K., Bridgewater, J., Gores, G. J. & Zhu, A. X. Systemic therapies for intrahepatic cholangiocarcinoma. J. Hepatol. 72, 353–363 (2020).

9. Doussot, A. et al. Outcomes after Resection of Intrahepatic Cholangiocarcinoma: External Validation and Comparison of Prognostic Models. J. Am. Coll. Surg. 221, 452–461 (2015).

10. Mavros, M. N., Economopoulos, K. P., Alexiou, V. G. & Pawlik, T. M. Treatment and Prognosis for Patients With Intrahepatic Cholangiocarcinoma: Systematic Review and Meta-analysis. JAMA Surg. 149, 565–574 (2014).

11. Hu, L.-S. et al. Recurrence Patterns and Timing Courses Following Curative-Intent Resection for Intrahepatic Cholangiocarcinoma. Ann. Surg. Oncol. 26, 2549–2557 (2019).

12. Ebata, T. et al. Randomized clinical trial of adjuvant gemcitabine chemotherapy versus observation in resected bile duct cancer. Br. J. Surg. 105, 192–202 (2018).

13. Edeline, J. et al. Gemcitabine and Oxaliplatin Chemotherapy or Surveillance in Resected Biliary Tract Cancer (PRODIGE 12-ACCORD 18-UNICANCER GI): A Randomized Phase III Study. J. Clin. Oncol. 37, 658–667 (2019).

14. Primrose, J. N. et al. Capecitabine compared with observation in resected biliary tract cancer (BILCAP): a randomised, controlled, multicentre, phase 3 study. Lancet Oncol. 20, 663–673 (2019).

15. Reames, B. N. et al. Impact of adjuvant chemotherapy on survival in patients with intrahepatic cholangiocarcinoma: a multi-institutional analysis. HPB 19, 901–909 (2017).

16. Akita, M. et al. Dichotomy in intrahepatic cholangiocarcinomas based on histologic similarities to hilar cholangiocarcinomas. Mod. Pathol. 30, 986–997 (2017).

17. Choi, W. J. et al. Systematic Review and Meta-Analysis of Prognostic Factors for Early Recurrence in Intrahepatic Cholangiocarcinoma After Curative-Intent Resection. Ann. Surg. Oncol. 29, 4337–4353 (2022).

18. AJCC Cancer Staging Manual. (2022).

19. Budau, K.-L. et al. Prognostic Impact of Tumor Budding in Intrahepatic Cholangiocellular Carcinoma. J. Cancer 13, 15 (2022).

20. Lowery, M. A. et al. Comprehensive Molecular Profiling of Intrahepatic and Extrahepatic Cholangiocarcinomas: Potential Targets for Intervention. Clin. Cancer Res. 24, 4154–4161 (2018).

21. Job, S. et al. Identification of Four Immune Subtypes Characterized by Distinct Composition and Functions of Tumor Microenvironment in Intrahepatic Cholangiocarcinoma. Hepatology 72, 965–981 (2020).

22. Martin-Serrano, M. A. et al. Novel microenvironment-based classification of intrahepatic cholangiocarcinoma with therapeutic implications. Gut gutjnl-2021–326514 (2022) doi:10.1136/gutjnl-2021-326514.

23. Jiao, Y. et al. Exome sequencing identifies frequent inactivating mutations in BAP1, ARID1A and PBRM1 in intrahepatic cholangiocarcinomas. Nat. Genet. 45, 1470–1473 (2013).

24. Jusakul, A. et al. Whole-Genome and Epigenomic Landscapes of Etiologically Distinct Subtypes of Cholangiocarcinoma. Cancer Discov. 7, 1116–1135 (2017).

25. Chaisaingmongkol, J. et al. Common Molecular Subtypes Among Asian Hepatocellular Carcinoma and Cholangiocarcinoma. Cancer Cell 32, 57–70.e3 (2017).

26. Dong, L. et al. Proteogenomic characterization identifies clinically relevant subgroups of intrahepatic cholangiocarcinoma. Cancer Cell 40, 70–87.e15 (2022).

27. Lin, J. et al. Multimodule characterization of immune subgroups in intrahepatic cholangiocarcinoma reveals distinct therapeutic vulnerabilities. J. Immunother. Cancer 10, e004892 (2022).

28. Cancer Genome Atlas Network. Comprehensive molecular portraits of human breast tumours. Nature 490, 61–70 (2012).

29. Gao, Q. et al. Integrated Proteogenomic Characterization of HBV-Related Hepatocellular Carcinoma. Cell 179, 561–577.e22 (2019).

30. Cao, L. et al. Proteogenomic characterization of pancreatic ductal adenocarcinoma. Cell 184, 5031–5052.e26 (2021).

31. Liu, Y., Beyer, A. & Aebersold, R. On the Dependency of Cellular Protein Levels on mRNA Abundance. Cell 165, 535–550 (2016).

32. Fröhlich, K. et al. Benchmarking of analysis strategies for data-independent acquisition proteomics using a large-scale dataset comprising inter-patient heterogeneity. Nat. Commun. 13, 2622 (2022).

33. Bosman, F. T., Carneiro, F., Hruban, R. H. & Theise, N. D. WHO classification of tumours of the digestive system. WHO Classif. Tumours Dig. Syst. (2010).

34. Föll, M. C. et al. Reproducible proteomics sample preparation for single FFPE tissue slices using acid-labile surfactant and direct trypsinization. Clin. Proteomics 15, 11 (2018).

35. Fiebig, H.-H. et al. Gene Signatures Developed from Patient Tumor Explants Grown in Nude Mice to Predict Tumor Response to 11 Cytotoxic Drugs. Cancer Genomics Proteomics 4, 197–209 (2007).

36. Ludwig, C. et al. Data-independent acquisition-based SWATH-MS for quantitative proteomics: a tutorial. Mol. Syst. Biol. 14, e8126 (2018).

37. Vidova, V. & Spacil, Z. A review on mass spectrometry-based quantitative proteomics: Targeted and data independent acquisition. Anal. Chim. Acta 964, 7–23 (2017).

38. Demichev, V., Messner, C. B., Vernardis, S. I., Lilley, K. S. & Ralser, M. DIA-NN: neural networks and interference correction enable deep proteome coverage in high throughput. Nat. Methods 17, 41–44 (2020).

39. Demichev, V. DIA-NN R package. (2020).

40. Nyamundanda, G., Poudel, P., Patil, Y. & Sadanandam, A. A Novel Statistical Method to Diagnose, Quantify and Correct Batch Effects in Genomic Studies. Sci. Rep. 7, 10849 (2017).

41. Rohart, F., Gautier, B., Singh, A. & Cao, K.-A. L. mixOmics: An R package for ‘omics feature selection and multiple data integration. PLOS Comput. Biol. 13, e1005752 (2017).

42. John, C. R. et al. M3C: Monte Carlo reference-based consensus clustering. Sci. Rep. 10, 1816 (2020).

43. Smyth, G. K. limma: Linear Models for Microarray Data. in Bioinformatics and Computational Biology Solutions Using R and Bioconductor (eds. Gentleman, R., Carey, V. J., Huber, W., Irizarry, R. A. & Dudoit, S.) 397–420 (Springer-Verlag, New York, 2005). doi:10.1007/0-387-29362-0_23.

44. Shannon, P. et al. Cytoscape: A Software Environment for Integrated Models of Biomolecular Interaction Networks. Genome Res. 13, 2498–2504 (2003).

45. Wu, T. et al. clusterProfiler 4.0: A universal enrichment tool for interpreting omics data. The Innovation 2, (2021).

46. Kanehisa, M. & Goto, S. KEGG: Kyoto Encyclopedia of Genes and Genomes. Nucleic Acids Res. 28, 27–30 (2000).

47. Kanehisa, M., Furumichi, M., Sato, Y., Kawashima, M. & Ishiguro-Watanabe, M. KEGG for taxonomy-based analysis of pathways and genomes. Nucleic Acids Res. 51, D587–D592 (2023).

48. Gillespie, M. et al. The reactome pathway knowledgebase 2022. Nucleic Acids Res. 50, D687–D692 (2022).

49. Ashburner, M. et al. Gene Ontology: tool for the unification of biology. Nat. Genet. 25, 25–29 (2000).

50. The Gene Ontology Consortium. The Gene Ontology resource: enriching a GOld mine. Nucleic Acids Res. 49, D325–D334 (2021).

51. Therneau, T. M. A Package for Survival Analysis in R. (2020).

52. Kassambara, A., Kosinski, M., Biecek, P. & Scheipl, F. survminer. (2021).

53. Cosenza-Contreras, M., et al. Fragterminomics: extracting information on proteolytic processing from shotgun proteomics data processed by FragPip. Preprint at 10.22541/au.169906623.39670856/v1 (2023).

54. Cao, J. et al. Intrahepatic Cholangiocarcinoma: Genomic Heterogeneity Between Eastern and Western Patients. JCO Precis. Oncol. 557–569 (2020) doi:10.1200/PO.18.00414.

55. Liberti, M. V. & Locasale, J. W. The Warburg Effect: How Does it Benefit Cancer Cells? Trends Biochem. Sci. 41, 211–218 (2016).

56. Mosele, F. et al. Recommendations for the use of next-generation sequencing (NGS) for patients with metastatic cancers: a report from the ESMO Precision Medicine Working Group. Ann. Oncol. Off. J. Eur. Soc. Med. Oncol. 31, 1491–1505 (2020).

57. Hyder, O. et al. Recurrence after operative management of intrahepatic cholangiocarcinoma. Surgery 153, 811–818 (2013).

58. Zhang, C. et al. Serum liver enzymes serve as prognostic factors in patients with intrahepatic cholangiocarcinoma. OncoTargets Ther. 10, 1441–1449 (2017).

59. Bian, J. et al. Characterization of Immunogenicity of Malignant Cells with Stemness in Intrahepatic Cholangiocarcinoma by Single-Cell RNA Sequencing. Stem Cells Int. 2022, 3558200 (2022).

60. Boonmars, T. et al. Involvement of c-Ski oncoprotein in carcinogenesis of cholangiocacinoma induced by Opisthorchis viverrini and N-nitrosodimethylamine. Pathol. Oncol. Res. POR 17, 219–227 (2011).

61. Kawamura, E. et al. Suppression of intrahepatic cholangiocarcinoma cell growth by SKI via upregulation of the CDK inhibitor p21. FEBS Open Bio 12, 2122–2135 (2022).

62. Shahinian, H., Tholen, S. & Schilling, O. Proteomic identification of protease cleavage sites: cell-biological and biomedical applications. Expert Rev. Proteomics 10, 421–433 (2013).

63. Fahrner, M., Kook, L., Fröhlich, K., Biniossek, M. L. & Schilling, O. A Systematic Evaluation of Semispecific Peptide Search Parameter Enables Identification of Previously Undescribed N-Terminal Peptides and Conserved Proteolytic Processing in Cancer Cell Lines. Proteomes 9, 26 (2021).

64. Fretwurst, T. et al. Proteomic profiling of human bone from different anatomical sites – A pilot study. PROTEOMICS – Clin. Appl. 16, 2100049 (2022).

65. Shahinian, J. H. et al. Proteomics highlights decrease of matricellular proteins in left ventricular assist device therapy†. Eur. J. Cardiothorac. Surg. 51, 1063–1071 (2017).

66. Lim, S. A. et al. Targeting a proteolytic neoepitope on CUB domain containing protein 1 (CDCP1) for RAS-driven cancers. J. Clin. Invest. 132, (2022).

67. Hidalgo, M. et al. Patient-Derived Xenograft Models: An Emerging Platform for Translational Cancer Research. Cancer Discov. 4, 998–1013 (2014).

