## Supplementary Figures for "Proteomic Characterization of Intrahepatic Cholangiocarcinoma Identifies Distinct Subgroups and Proteins Associated with Time-To-Recurrence"

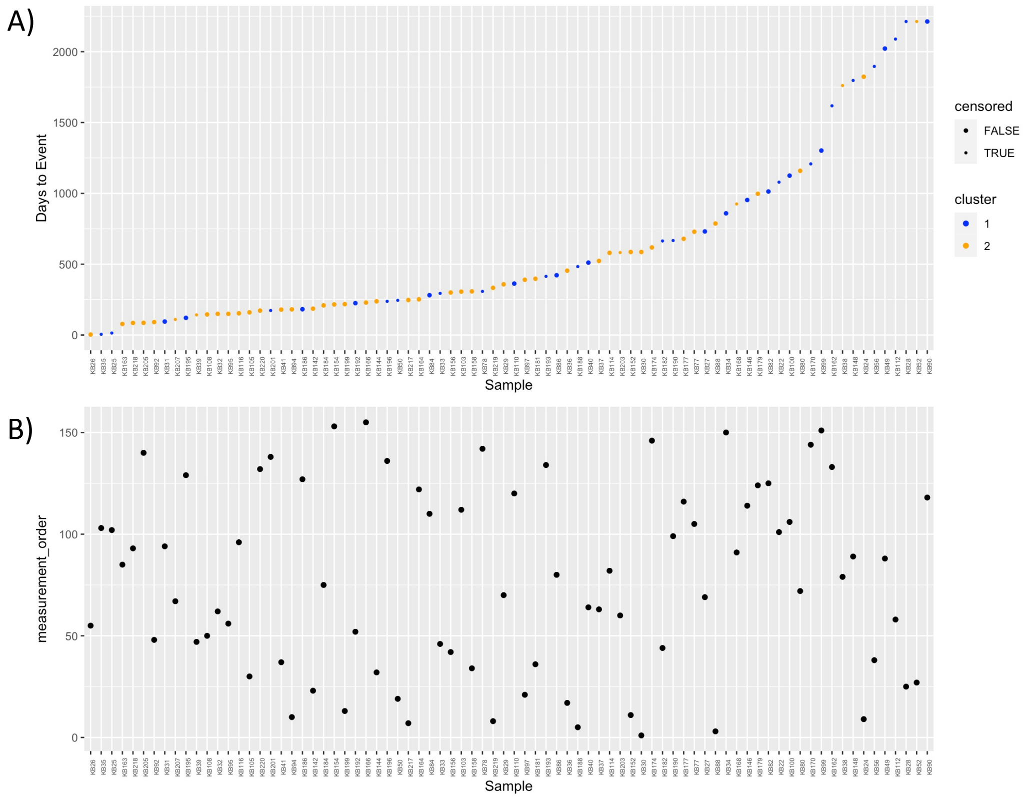


***Figure S1 - TTR distribution across cohort:*** *A) TTR distribution of cohort. B) Matching measurement order of samples.*

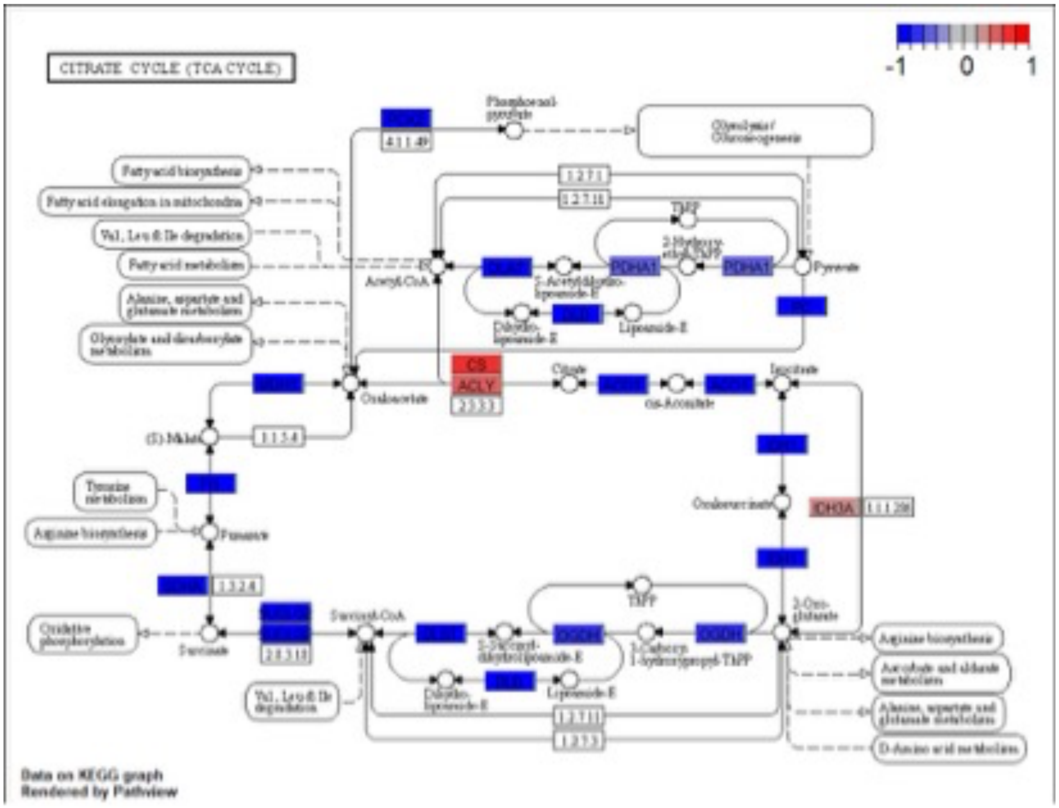
CITRATE CYCLE (TCA CYCLE)
-1 0 1
Phosphoenol-pyruvate
4.1.1.49
Glycolysis / Gluconeogenesis
Fatty acid biosynthesis
Fatty acid elongation in mitochondria
Val, Leu & Ile degradation
Fatty acid metabolism
Alanine, aspartate and glutamate metabolism
Glyoxylate and dicarboxylate metabolism
1.2.7.1
1.2.7.11
TPP
PDHA1
Acetyl-CoA
Pyruvate
Lipoamide-E
CS
ACLY
Citrate
cis-Aconitate
Isocitrate
Oxaloacetate
(S)-Malate
Tyrosine metabolism
Arginine biosynthesis
Fumarate
Oxalosuccinate
IDH3A
Succinyl-CoA
OGDH
2-Oxo-glutarate
Oxidative phosphorylation
Succinate
Arginine biosynthesis
Ascorbate and aldarate metabolism
Alanine, aspartate and glutamate metabolism
D-Amino acid metabolism
Data on KEGG graph
Rendered by Pathview

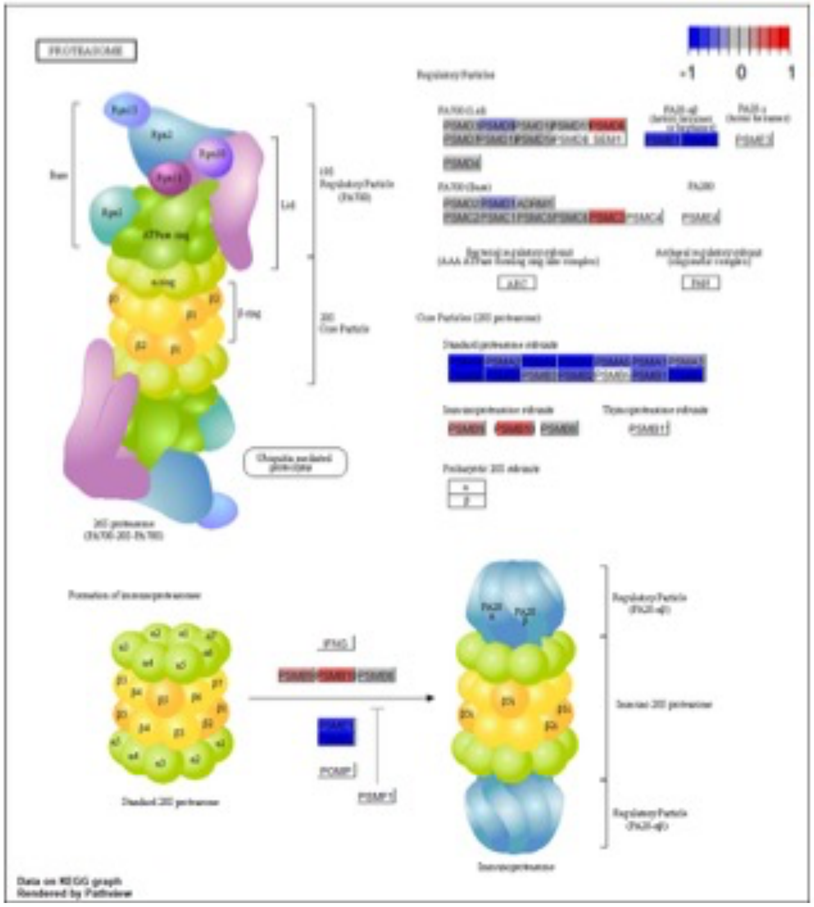
PROTEASOME
-1 0 1
Data on KEGG graph
Rendered by Pathview

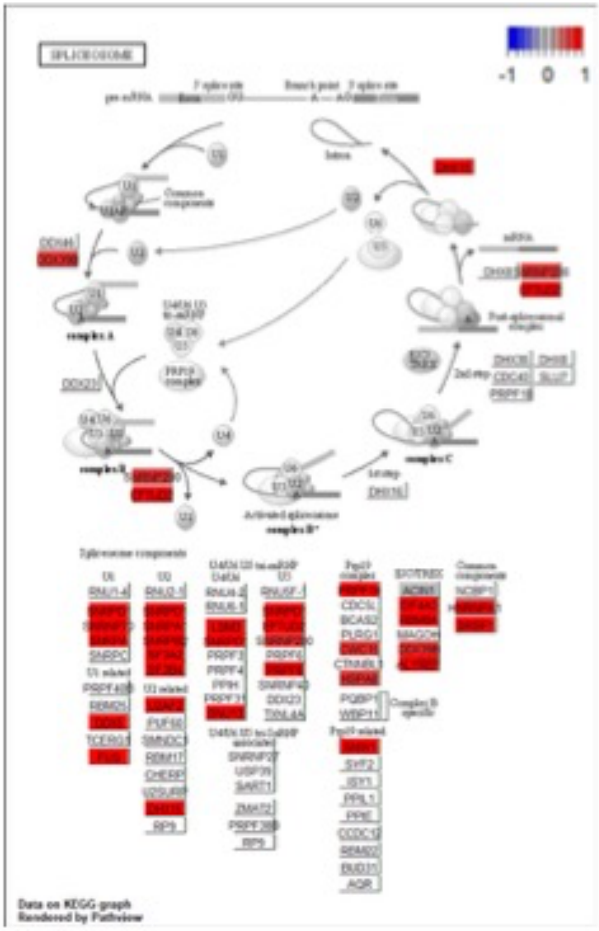
SPLICEOSOME
-1 0 1
Data on KEGG graph
Rendered by Pathview

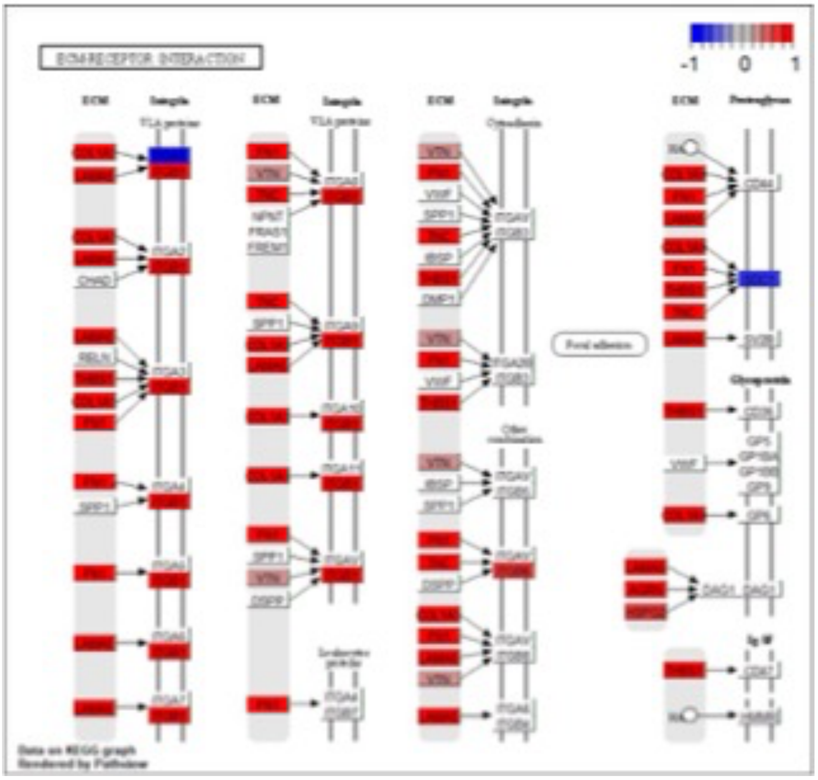
ECM-RECEPTOR INTERACTION
-1 0 1
Data on KEGG graph
Rendered by Pathview

***Figure S2 - KEGG enrichment Tumor vs. TANM:*** *Tumor in red, TANM in blue. Selected KEGG enrichment results of Limma tumor vs. TANM.*

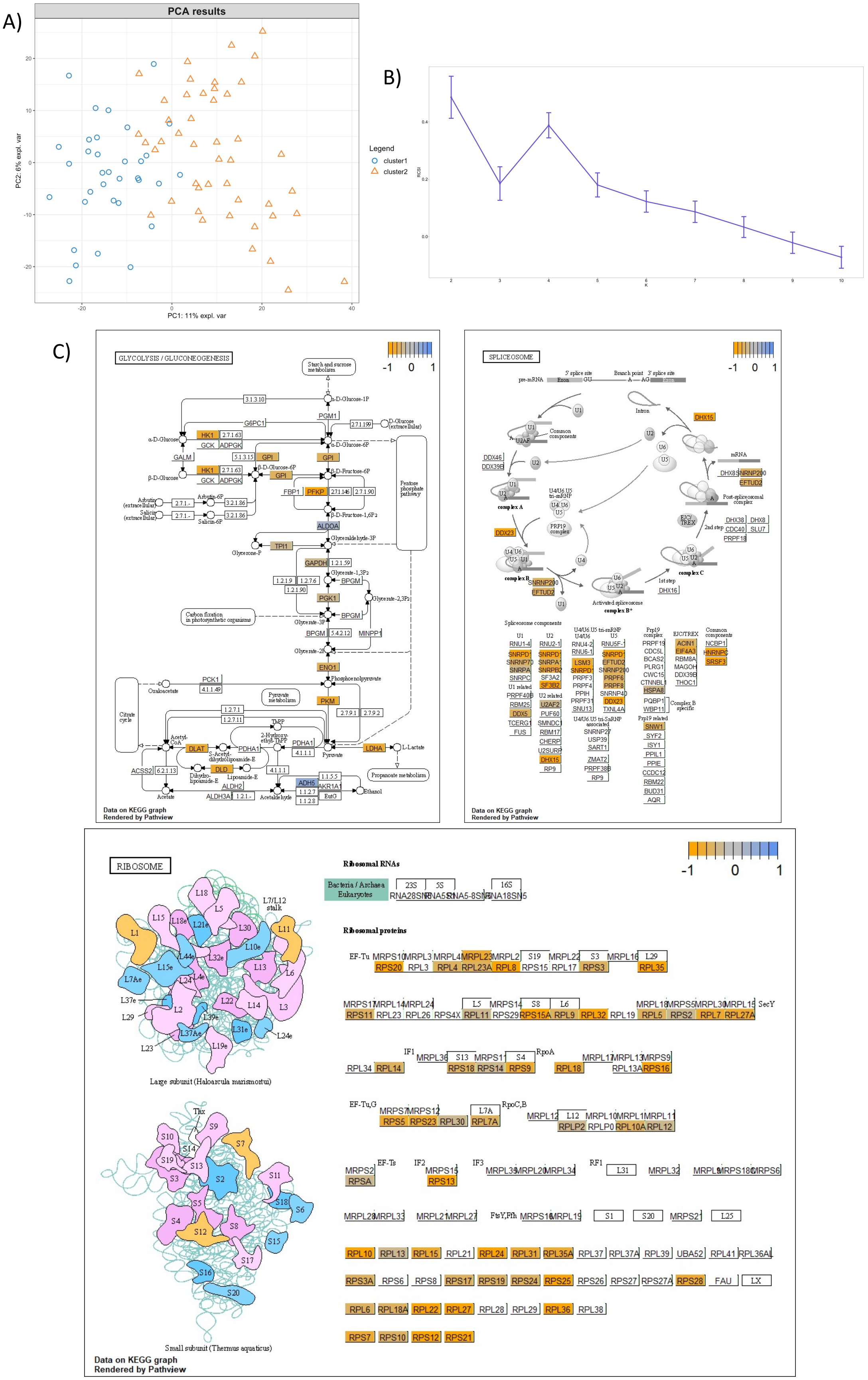

A)
PCA results
PC1: 11% expl. var
PC2: 6% expl. var
Legend
cluster1
cluster2
B)
C)
GLYCOLYSIS / GLUCONEOGENESIS
SPLICEOSOME
RIBOSOME
-1 0 1
Ribosomal RNAs
Ribosomal proteins
Large subunit (Haloarcula marismortui)
Small subunit (Thermus aquaticus)
Data on KEGG graph
Rendered by Pathview

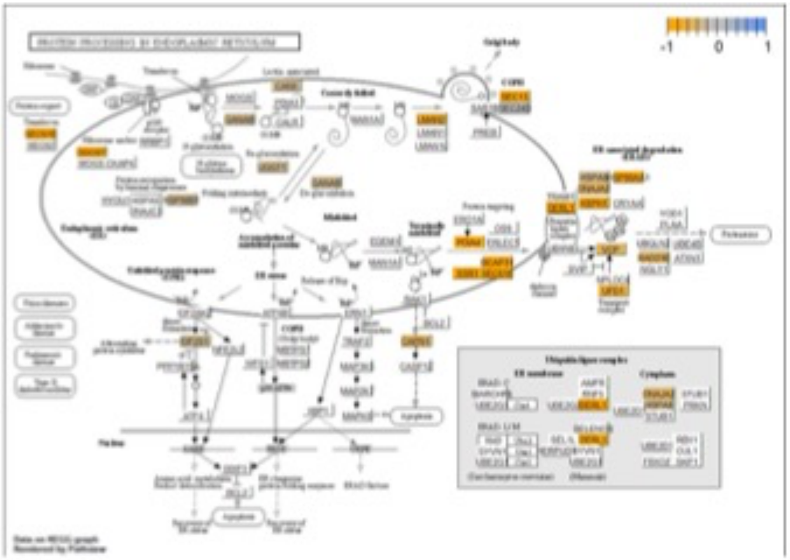
-1 0 1
PROTEIN PROCESSING IN ENDOPLASMIC RETICULUM
Data on KEGG graph
Rendered by PathVisio

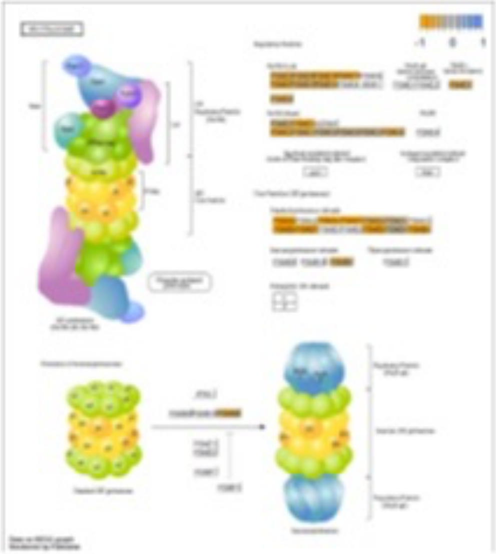
-1 0 1

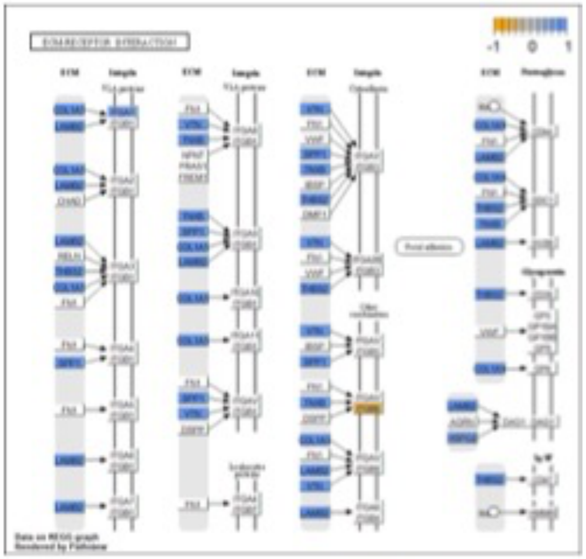
-1 0 1
ECM-RECEPTOR INTERACTION
Data on KEGG graph
Rendered by PathVisio

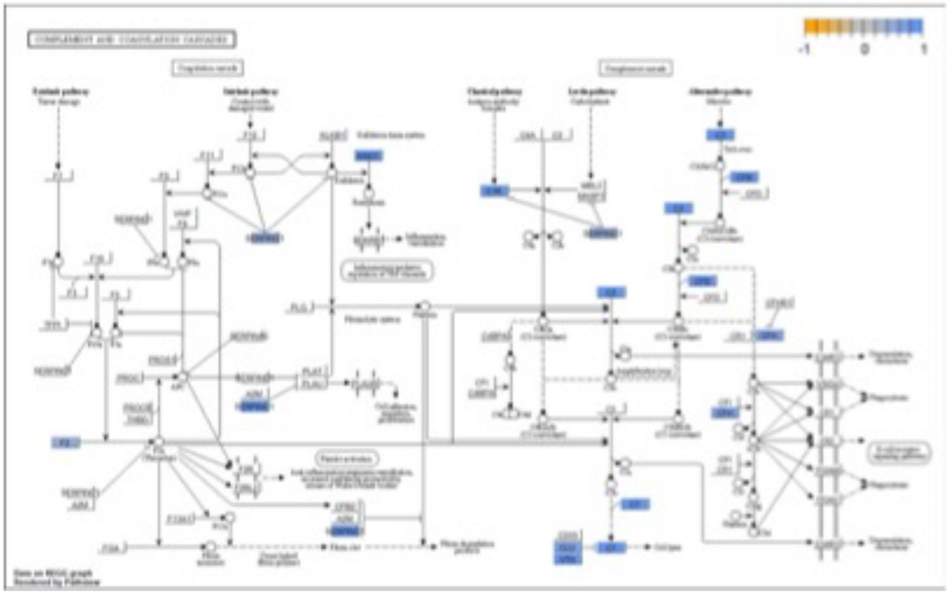
-1 0 1
COMPLEMENT AND COAGULATION CASCADES
Data on KEGG graph
Rendered by PathVisio

***Figure S3 - KEGG enrichment cluster 1 vs. cluster 2:*** *Cluster 1 in blue, cluster 2 in orange. A) PCA of tumor samples. B) Monte-Carlo simulation result of clustered tumor samples. Potential numbers of subclusters k can be selected through the Relative Cluster Stability Index (RCSI).* ***C)*** *Selected KEGG enrichment results of Limma cluster 1 vs. cluster 2.*

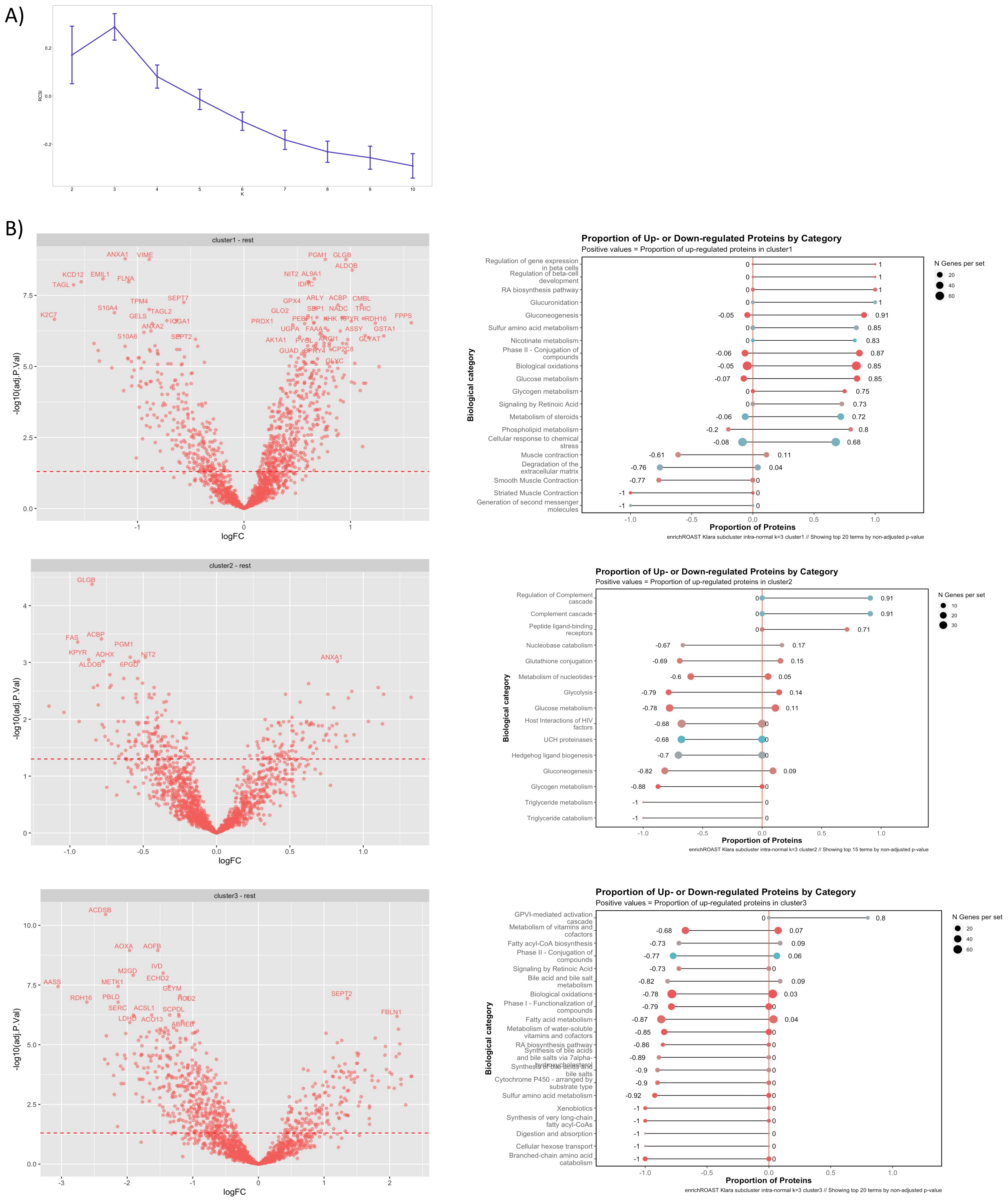


***Figure S4 - TANM clusters:*** *TANM tissue clustering. A) Monte-Carlo simulation result of clustered TANM samples. Potential numbers of subclusters k can be selected through the Relative Cluster Stability Index (RCSI). B) Volcano plots of identified TANM clusters and matching enrichment results.*

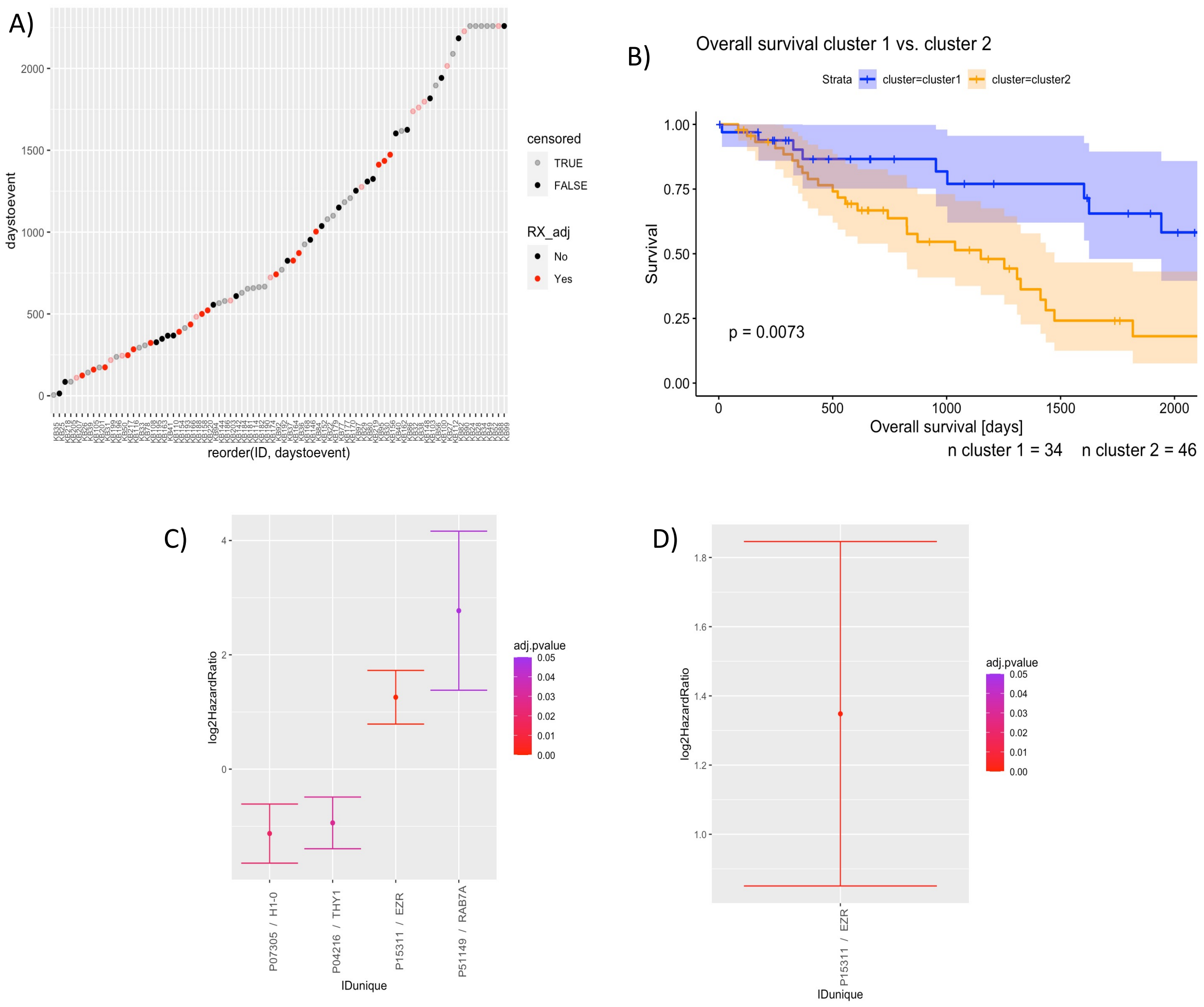


***Figure S5 - CPHM overall survival:*** *Statistical analysis of overall survival data. A) Overall survival distribution across cohort. B) Kaplan-Meier plot of overall survival distribution between clusters. C) Result of CPHM performed with overall survival data without covariates. D) Result of CPHM performed with overall survival data with radiotherapy as covariate.*
